# Direct reprogramming of adult hepatocytes to generate LGR5+ endodermal progenitor

**DOI:** 10.1101/2020.05.29.123273

**Authors:** Diana Chaker, Christophe Desterke, Nicolas Moniaux, Tony Ernault, Noufissa Oudrhiri, Jamila Faivre, Ali G Turhan, Annelise Bennaceur-Griscelli, Frank Griscelli

## Abstract

We successfully converted hepatocytes isolated from adult mice into expandable and stable leucine-rich repeat-containing G-protein-coupled receptor 5 (LGR5)-positive endodermal progenitor cells (EndoPCs). This was accomplished *in vitro* through transient exposure to four transcriptional factors (OCT3/4, SOX2, KLF4, and cMYC) and STAT3 activators. EndoPCs were generated by a process that involved an epithelial-mesenchymal transition (EMT) without the use of any components of the canonical WNT/β-catenin or LGR5/R-spondin signaling pathways. We showed that the proliferation and capacity for self-renewal of EndoPCs in 2D long-term culture were controlled by three interrelated signaling pathways: gp130/JAK/STAT3, LGR5/R-spondin, and WNT/β-catenin. After long-term maintenance in two- and three-dimensional culture systems, EndoPCs were able to differentiate into liver-restricted lineages such as hepatocyte-like cells and bile duct-like structures *in vitro* and *in vivo*. After intra-muscular injection, EndoPCs generated macroscopically visible and well-vascularized liver-like tissue, which contained Alb^+^ liver parenchyma-like structures and substantial, KRT7/KRT19^+^ bile duct-like cell organizations.

**Conclusion:** We have developed an efficient method for producing LGR5^+^ adult endodermal stem cells. These cells will be useful for the *in vitro* study of the molecular mechanisms of liver development and have important potential for therapeutic strategies, including approaches based on bioengineered liver tissue. These cells also open up new avenues for experiments focused on disease modeling, toxicology studies, and regenerative medicine applications.

## INTRODUCTION

The liver has a significant latent capacity for regeneration. After extreme stress or chronic injury, a population of atypical ductal bipotent progenitor cells (BPCs) emerges from the bile ducts(*1, 2*) and is able to differentiate into both hepatocytes and biliary cells, as demonstrated by lineage tracing after liver injury(*3, 4*). However, liver BPCs and the signaling pathways that maintain the progenitor fate within the liver have not yet been fully characterized. In contradiction of conventional BPC theory, several groups have demonstrated that hepatocytes can be efficiently reprogrammed into proliferative BPCs in response to chronic liver injury (*5–8*). Recently, a technique based on the use of three small chemical molecules (Y-27632, A-83-01, CHIR99021) was demonstrated to successfully convert mature hepatocytes from rats and mice into proliferative liver progenitor cells; these cells were shown to differentiate into both mature hepatocytes and biliary epithelial cells, and were able to repopulate chronically injured liver tissue in transgenic mice that carried the urokinase-type plasminogen activator (uPA) gene(*9*). *In vivo*, these chemically induced liver progenitors were shown to preferentially give rise to hepatocytes, and more rarely to bile duct structures. However, these reports did not clearly identify the signaling pathways involved either in *in vivo* differentiation or in *in vitro* maintenance of these cell populations in culture. In addition, these biliary epithelial cells were shown to express only KRT19, which is a hallmark of non-differentiated ductal bipotent progenitor cells and liver cancer cells(*10*).

The leucine-rich repeat-containing G-protein-coupled receptor 5 (LGR5) protein was recently shown to be a marker of adult stem cells in the murine liver,(*3*) which have been grown *in vitro* exclusively in the context of organoids in a three-dimensional long-term culture system that utilizes high doses of WNT/β-catenin and LGR5 ligands. Human LGR5^+^ liver stem cells that expressed both ductal (KRT19, SOX9, OC2) and hepatocyte markers (HNF4α) were also recently isolated and expanded into 3D organoid structures that were efficiently converted into functional hepatocytes *in vitro* and upon transplantation *in vivo*(*11*).

Here we show for the first time that mature hepatocytes can be directly converted *in vitro* into LGR5^+^ proliferative endodermal progenitor cells (EndoPCs) after transient exposure to four embryonic transcriptional factors, OCT4, SOX2, KLF4, and c-MYC, and a high dose of STAT3 activators. We demonstrate that EndoPCs are dependent on STAT3 activators to regulate gp130/JAK/STAT3, as well as on the LGR5/R-spondin and WNT/β-catenin signaling pathways, which enable their proliferation and their capacity for self-renewal in 2D long-term culture conditions. In addition, EndoPCs express typical endodermic transcriptional factors involved in endodermic lineage development thus permitting them to differentiate easily *in vitro*, and spontaneously *in vivo*, into both functional hepatocytes and biliary cells.

## RESULTS

### Establishment and characterization of endodermal progenitor cells

In the presence of mLIF, 10^6^ murine hepatocytes were transduced with 20 MOI of recombinant adenoviruses that harbored human OCT4, SOX2, KLF4, and cMYC transgenes. This process resulted in the emergence of more than 20 different expandable (Fig. S1) and genetically stable clones (Fig. S2). Non-infected control hepatocytes failed to survive a second passage, but all transduced clones rapidly proliferated. After 10 iterative passages, these cells exhibited a typically mesenchymal phenotype, instead of the epithelial morphology of iPSCs (Fig. S1E). We thus hypothesized that the clones were partially reprogrammed cells which had become ‘stuck’ during reprogramming and were unable to complete the pluripotency program. Instead of generating iPSCs, we presumed that our protocol had produced endodermal progenitor cells (EndoPCs). This hypothesis was supported by the results of the flow cytometry analysis, which revealed the absence of stage-specific embryonic antigen (SSEA)-1 expression (Fig. 1A, Table S1), and the presence of mesenchymal markers such as CD29, CD73, CD51, CD166, and Sca-1 (Fig. S3, Table S1).

**Figure 1:**
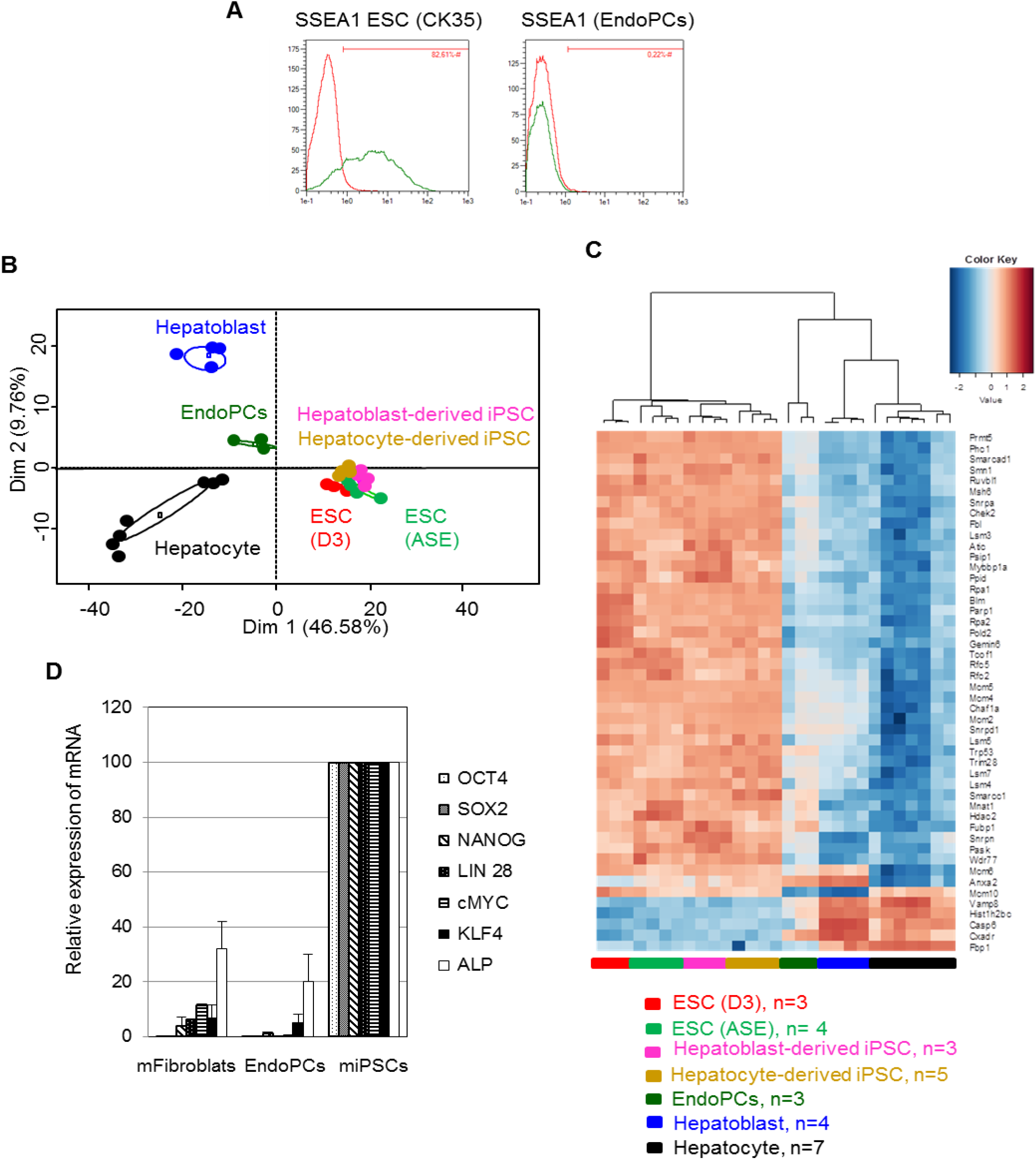
EndoPC characterization. (**A**) Flow cytometry analysis of pluripotency marker SSEA-1 in murine embryonic stem cells (ESC/CK35) and EndoPCs. (**B**) Unsupervised principal component analysis performed on 299 Plurinet genes from transcriptome analysis of EndoPCs (dataset GSE51782) compared to murine embryonic stem cells (ESCs/D3 and ASE), murine hepatocyte-derived induced pluripotent stem cells (iPSCs), hepatoblast-derived iPSCs, hepatocytes, and hepatoblasts isolated from a pool of fetal (E16.5) livers (GSE33110). All samples were produced in and isolated from C57BL/6 mice. On each axis, the percentage of variance is detailed. Ellipses and shapes represent clustering patterns of the samples. (**C**) Gene expression heatmap and unsupervised classification based on expression changes of Plurinet genes in ESCs, iPSCs, EndoPCs, hepatoblasts, and hepatocytes (Pearson distances). (**D**) Differences in expression, as revealed by quantitative RT-PCR, of genes that are characteristic of iPSCs among EndoPCs, murine iPSCs, and primary murine C57BL/6 fibroblasts. EndoPCs were tested at passage 30, iPSCs at passage 10, and fibroblasts at passage 3. Seven different factors— OCT4, SOX2, NANOG, LIN28, CMYC, KLF4, and alkaline phosphatase (ALP)—were quantified and subsequently normalized to the mRNA level found in iPSCs (value of 100).

To confirm the flow cytometry results, we profiled the overall gene expression of three different EndoPC clones and analyzed these data with respect to the Plurinet genes, as described by Müller et al.(*12*) To facilitate the comparison between EndoPCs and different cell populations, we assembled a dataset of expression data from adult hepatocytes, fetal hepatoblasts, and different sources of murine pluripotent stem cells, including murine embryonic stem cells (ESCs) (D3 or ASE) and murine iPSCs that were derived from either hepatocytes or fetal hepatoblasts. This comparison revealed that EndoPCs were mathematically midway between pluripotent stem cells (iPSCs and ESCs) and the hepatocytes that had been used to derive the EndoPCs (Fig. 1B,C), providing further evidence that EndoPCs were not themselves pluripotent. These results were confirmed by a qRT-PCR analysis that failed to detect iPSC-enriched genes such as *OCT4, SOX2, NANOG, LIN28, CMYC, KLF4*, and alkaline phosphatase (*ALP*) in EndoPC samples, in contrast to results from miPSCs and murine fibroblasts (Fig. 1D).

We next investigated the endodermal differentiation stage of EndoPCs by integrating our transcriptome data with a dataset comprised of murine samples (GSE13040) that included ESCs (D3) and samples isolated from E8.5 and E11.5 fetal livers. An unsupervised principal component analysis based on the Plurinet signature was able to significantly (p=0.00006) distinguish among samples on the first principal axis based on their stage of endodermal differentiation; EndoPCs were relatively distant from ESCs (D3) and E8.5 samples, and were instead closer to E11.5 liver samples (Fig. S4A). Likewise, an unsupervised clustering revealed that EndoPCs were more similar to the sample isolated from E11.5 fetal livers. In both groups, the expression of pluripotent genes was repressed (Fig. S4B), showing that EndoPCs represent a new source of primitive fetal liver stem cells. This relationship was also confirmed with an analysis of mRNA (Fig. S5A) and by immunofluorescence staining (Supporting Fig. 5B) that demonstrated that EndoPCs correctly expressed SOX17, CXCR4, GATA4, and the Foxa subfamily, which are transcriptional factors involved in early liver development. EndoPC clones also expressed transcriptional factors involved in the development of bile ducts (*KRT7/19, Hnf6*, *Hnf1β*) and hepatocytes (*KRT8/18, Hnf4α, Hnf1α*, *AFP, ALB*) (Fig. S6, Table **S**1); these factors have been implicated in liver ontology, which suggested that EndoPCs could have potential for commitment toward both cholangiocytes and hepatocytes.

### EndoPCs exhibit a LGR5^+^ liver stem cell signature

Since LGR5 was recently shown to be a marker of adult murine liver stem cells,(*4*) we investigated whether gene expression in EndoPCs contained a signature that could distinguish these cells as liver stem cell progenitors. We first compared the gene expression profile of LGR5^+^-sorted liver cells to primary hepatocytes using published datasets (GSE32210).(*4*) Using a supervised analysis performed on both samples with the significance analysis for microarray (SAM) algorithm, we were able to identify 275 different gene probes that were upregulated in LGR5^+^ cells compared to hepatocytes (more than two-fold change and a false discovery rate less than 0.05; Fig.1A,B, Table S2). The expression patterns of these upregulated genes in LGR5^+^ cells also enabled discrimination among transcriptome samples using unsupervised classification (heatmap with Euclidean distances, Fig. 2A and Table S2).

**Figure 2:**
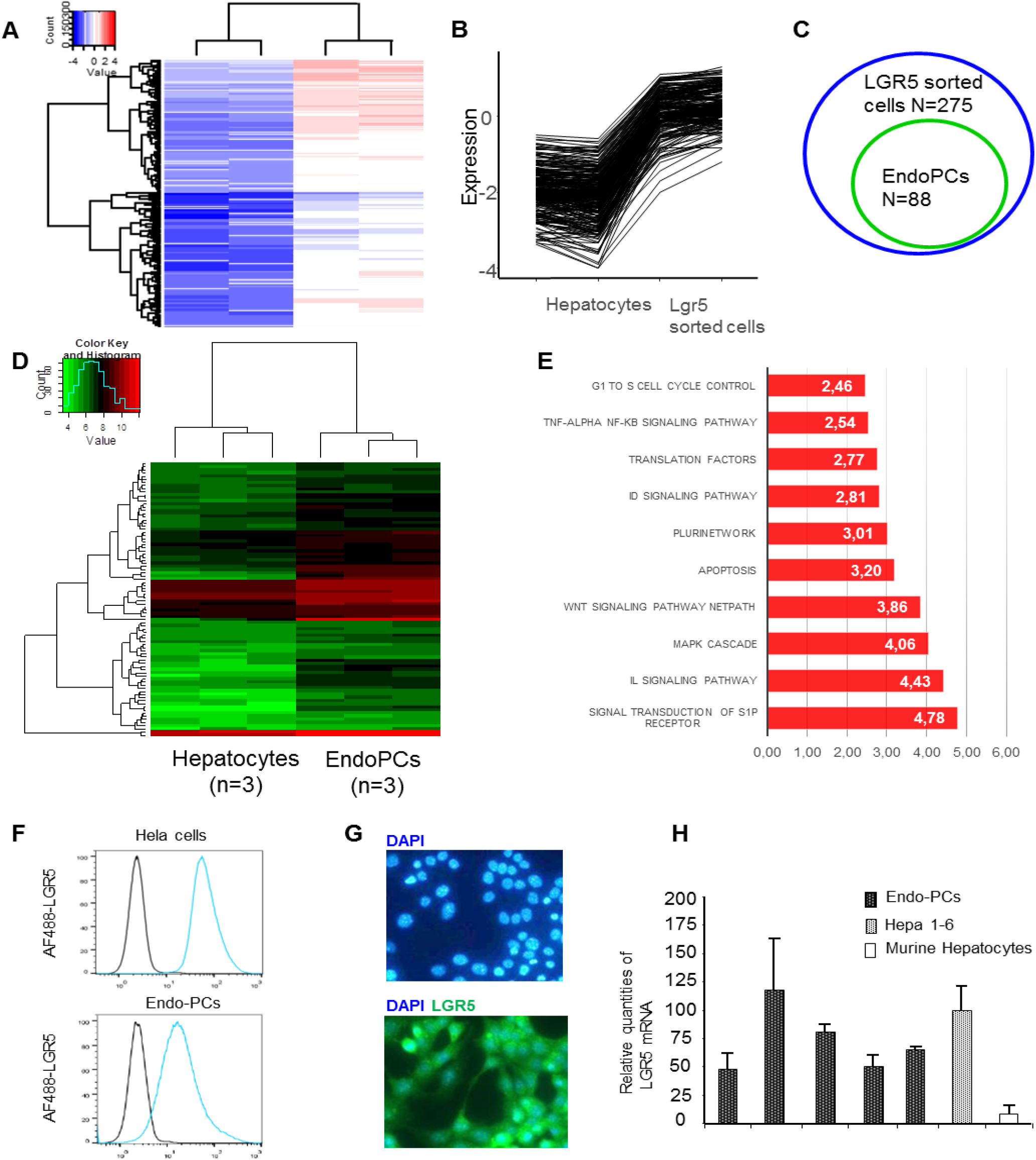
Gene expression profile of LGR5^+^ EndoPCs by microarray. (**A**) Unsupervised classification of the expression profiles of sorted LGR5^+^ liver cells compared to hepatocytes, from published datasets (GSE32210). (**B**) Cord plot of gene expression, indicating genes that were upregulated in LGR5^+^ cells compared to hepatocytes. (**C**) Venn diagram of gene overlap between genes that were upregulated in LGR5^+^ cells and in EndoPCs. (**D**) Expression heatmap of genes that were upregulated in both EndoPCs and LGR5^+^ cells. (**E**) Barplot of Z-scores obtained from an analysis of functional enrichment (using the Wikipathway database) of LGR5-dependent genes that were overexpressed in EndoPCs. (**F**) Expression of LGR5 in Hela cells and EndoPCs as determined by flow cytometry analysis. (**G**) Immunofluorescence staining showing the expression of LGR5 in EndoPCs. DNA was stained with DAPI. (**H**) Results of quantitative RT-PCR for the detection of LGR5 mRNA in five individual EndoPC clones, in Hepa 1-6, and in murine hepatocytes.

Compared to the expression profiles of primary hepatocytes, we identified 88 genes that were upregulated in LGR5^+^ cells. This signature was also able to discriminate between LGR5^+^ liver cells and EndoPCs using an unsupervised classification (Venn Diagram, Fig. 2C, heatmap with Euclidean distances, Fig.,2D, and Table S3). These 88 LGR5-related genes were subjected to an analysis of functional enrichment using the Wikipathway database, which revealed an association with several pathways that regulate stem cell proliferation; these included the WNT pathway, a set of genes belonging to the PluriNetWork, and the TNFα NF-KB signaling pathway (Fig. 2E). Because the presence of LGR5 in the EndoPCs was verified by flow cytometry (Fig. 2F, Table S1), immune-fluorescence staining (Fig. 2G) and by RT-PCR (Fig. 2H), we thus concluded that EndoPCs represent a new source of LGR5^+^ endodermal stem-like cells.

### EndoPC expansion and self-renewal depends on gp130/JAK/STAT3, LGR5/R-spondin, and WNT/β-catenin signaling pathways

All EndoPC clones continuously expanded in the presence of STAT3 activators (mLIF or mIL6), and we were able to passage them over 50 times in culture, which suggested that EndoPCs have a capacity for self-renewal. To verify whether the gp130/JAK/STAT3 pathway plays a role in this ability, EndoPCs were maintained for 7 days with low-serum-containing medium with or without mLIF. The withdrawal of mLIF significantly inhibited their proliferation (Fig. 3A) but without apparent mortality (Fig. S7). The addition of a pan-JAK inhibitor into the mLIF-containing culture medium was followed by a significant inhibition of EndoPC proliferation, similar to what was observed in the culture without mLIF (Fig. 3A). These results confirmed that the gp130/JAK/STAT3 pathway is involved in the proliferation and self-renewal of EndoPCs. As expected, mLIF could be substituted with 100 ng/ml of IL6, which likewise enabled long-term proliferation (Fig. S8) and activated the gp130/JAK/STAT3 pathway, as revealed by the detection of the Tyr-705-phospho-STAT3 form by western blot analysis (Fig. 3B)

**Figure 3:**
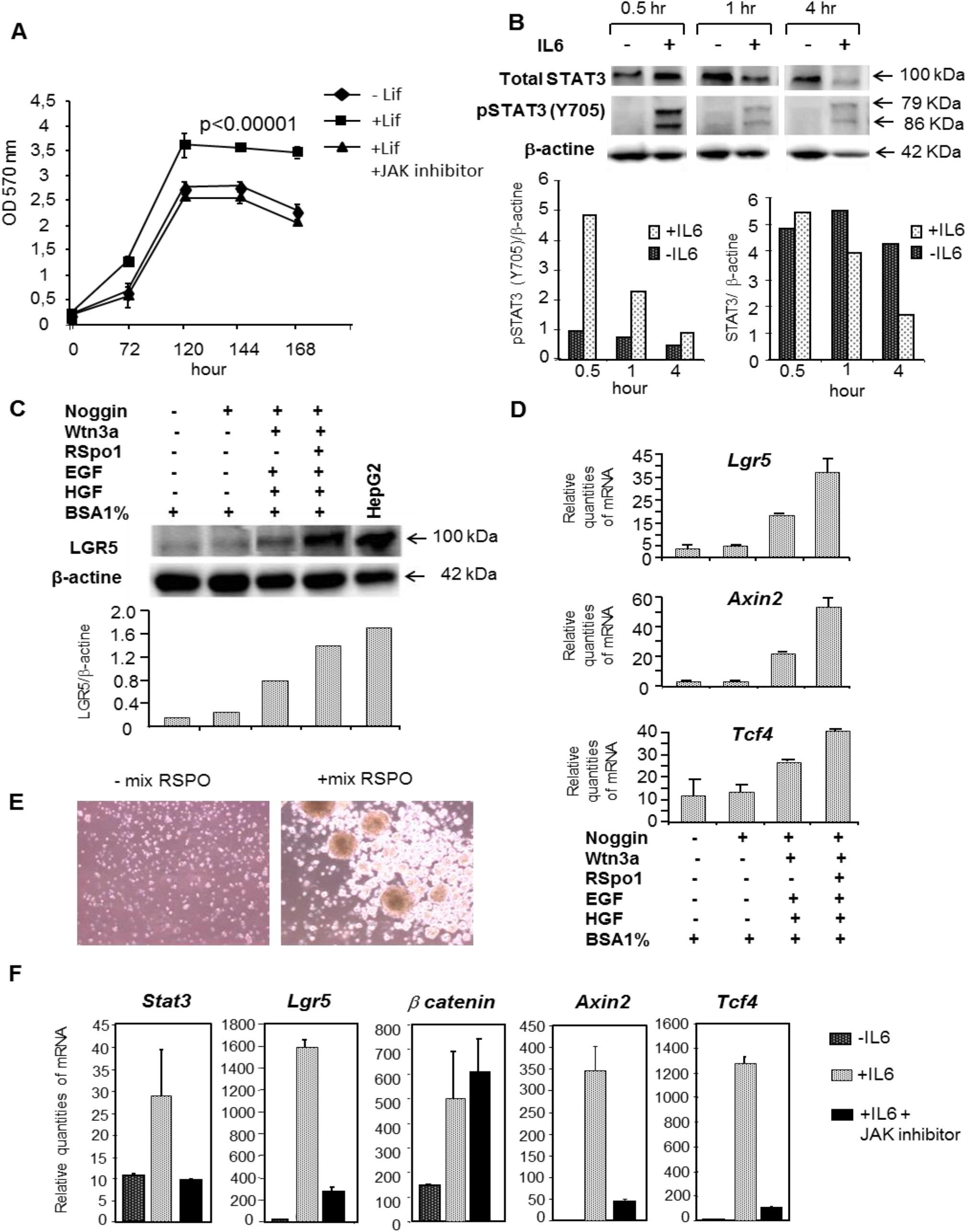
EndoPCs are dependent on both gp130/JAK/STAT3 and WNT/β-catenin signaling pathways. (**A**) Proliferation Assay (MTT Assay) performed on EndoPCs after 72, 120, 144, or 168 hours in culture with or without 1000 units/ml of LIF, compared to the condition with mLIF and 5 μM of pan-JAK inhibitor. (**B**) Western blot analysis of STAT3, pSTAT3 (Y705), and β-actin in EndoPCs that expanded with or without 100 ng/ml of IL6. Analysis was performed after 0.5, 1, and 4 hr of exposure; all bands were quantified relative to the intensity of β-actin. (**C**) Western blot analysis of LGR5 and β-actin in EndoPCs that expanded in the presence or absence of five different cytokines (RSPO mix, containing 1% BSA, 30 ng/ml of mWNT3a, 500 ng/ml of mRspo1, 20 ng/ml of mEGF, 50 ng/ml of mHGF, and 50 ng/ml of mNoggin). All bands were quantified relative to the intensity of β-actin. (**D**) Results of quantitative RT-PCR used to detect *Lgr5*, *Axin2*, and *Tcf4* in EndoPCs that expanded in the presence or absence of RSPO mix. (E) Organoid-like structures in low-attachment plates that developed in the presence or absence of RSPO mix. (**F**) Results of quantitative RT-PCR used to detect *Stat3*, *Lgr5*, *βcatenin*, *Axin2*, and *Tcf4* in EndoPCs that expanded for 8 hr with or without 100 ng/ml of IL6, compared to EndoPCs expanded with 100 ng/ml IL6 and 5 μM of pan-JAK inhibitor.

EndoPCs were also shown to be responsive to both the LGR5/R-spondin and WNT/β-catenin signaling pathways. This was demonstrated through the addition of five agonists of both pathways to a serum-free medium; the final composition of the medium was 1% BSA, 30 ng/ml mWNT3a, 500 ng/ml mRspo1, 20 ng/ml mEGF, 50 ng/ml mHGF, and 50 ng/ml mNoggin (RSPO mix). EndoPCs that were cultured with the RSPO mix demonstrated significantly increased proliferation compared to cells which lacked this mix (Supporting Fig. 9); the presence of the RSPO mix was also associated with the upregulation of LGR5 protein as determined by western blot analysis (Fig. 3C), and with the expression of *LGR5*, *Axin2*, and *Tcf4* mRNA (Fig. 2D). Furthermore, EndoPCs that were exposed to the RSPO mix for 7 days in low-attachment plates generated organoid-like structures (Fig. 2E). The proliferation of EndoPCs that was stimulated by the RSPO mix was downregulated by a JAK inhibitor (Fig. S8), which we interpreted as evidence of cross-talk between the gp130/JAK2/STAT3 and WNT/β-catenin pathways. To confirm this, we investigated whether activation of the gp130/JAK/STAT3 pathway could modify the behavior of the WNT/β-catenin pathway. We found that gp130/JAK/STAT3 signaling could efficiently activate the WNT/β-catenin signaling pathway by controlling the expression of *Lgr5*, *Axin2*, *Tcf4*, and *β catenin* (Fig. 3F). Indeed, the addition of IL6 activated the gp130/JAK/STAT3 pathway and resulted in a drastic increase in the number of transcripts of *LGR5*, *Axin2*, and *Tcf4*. Instead, these effects were abolished by the addition of a JAK inhibitor to the culture medium (Fig. 3F).

### Differentiation of EndoPCs into both hepatocytes and bile duct structure in vivo

To verify that EndoPCs were able to spontaneously differentiate *in vivo* in a non-hepatic microenvironment, 3×10^6^ EndoPCs were injected intra-muscularly. Macroscopically visible (523+50 mm^3^, n=3) and well-vascularized liver-like tissues developed after 49–101 days (Fig. 4A). H&E staining revealed two main types of tissues: areas of bile duct-like organization (Fig. 4B), which expressed bile duct markers (KRT19, KRT7, and acid mucosubstances), and areas of liver parenchyma-like structures (Fig. 4C), which expressed hepatocyte markers (Alb, glycogen). The liver tissue was highly vascularized (Fig. 4D), with a strong connection to pre-existing blood vessels that enabled the liver to grow and survive for several weeks. Using RT-PCR, we confirmed the commitment of EndoPCs into both ductal (positive for *KRT7/19*) and hepatic functional structures (positive for *KRT8/18*, *Alb*, *G6P, Cyp2E1, Cyp3A11, Cyp2D22, Cyp2C55*; Fig. 4E). To further characterize the activity of Cyp3A11 in these tissues, we used mass spectrometry to investigate the pharmacokinetics of midazolam, which is metabolized in the liver by Cyp3A11 to 1-hydroxymidazolam by glucuronidation. We analyzed tissue extracts from two mice and compared them to extracts from a murine mESC-derived teratoma; in doing so, we observed a five-fold increase in 1-hydroxymidazolam in EndoPC-derived teratoma compared to the control extracts. This result pointed to the existence of functional Cyp3A11 within both liver-like tissues (Fig. 4F).

**Figure 4:**
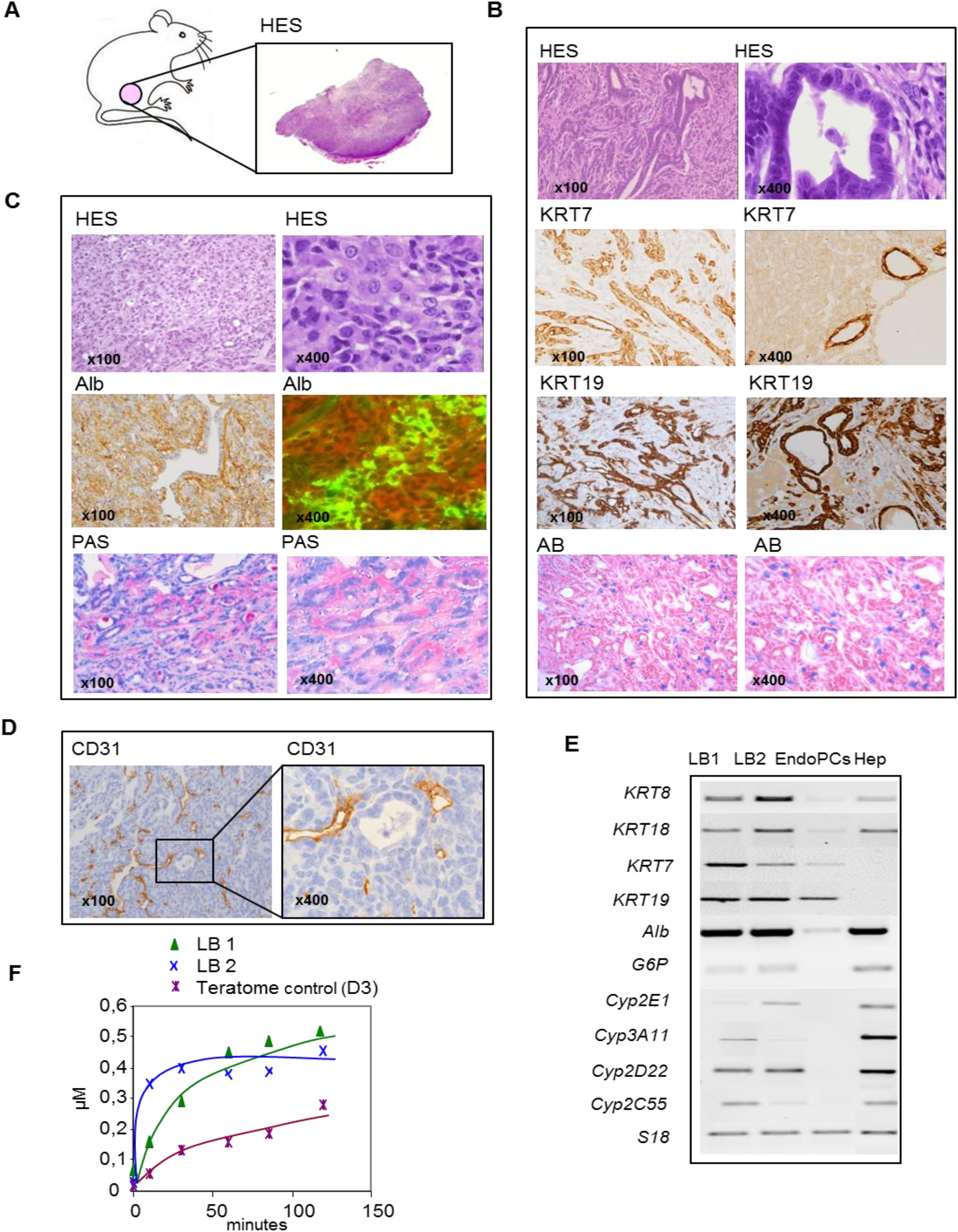
Intramuscular injection of EndoPCs generates liver-like structures. (**A**) Picture of a mouse with an emerging liver-like tissue, 3 months following intramuscular injection of 5×10^6^ EndoPCs into C57BL/6 mice. (**B**) Alcian Blue (AB) staining reveals branching-like structures that were immunoreactive for KRT7 and KRT19, as well as acid mucosubstances and polysaccharides. (**C**) Immuno-fluorescence and immuno-histochemical analysis revealed that the liver-like structures contained parenchyma-like structures positive for Albumin (Alb) as well as large areas of glycogen (revealed by periodic acid-Schiff staining). (**D**) Large areas of blood vessels were positive for PECAM1 (CD31). (E) RT-PCR analyses of hepatic markers in two different liver-like tissues (LB1 and LB2) compared to EndoPCs and murine hepatocytes (Hep). (**F**) The pharmacokinetics of midazolam and its main metabolite, 1-hydroxymidazolam, as quantified by mass spectrometry of extracts from EndoPC-derived liver-like tissues (LB1 and LB2) and ESC (D3)-derived teratoma.

In order to document the potential of EndoPCs to colonize the liver parenchyma, we injected either PBS or 3×10^6^ cells of EndoPC-luc, which expressed the luciferase reporter gene, into the spleens of five adult C57BL/6 mice that had undergone 30% partial hepatectomy. After 7 days, firefly bioluminescence imaging detected luciferase activity exclusively in the liver and the spleen of all five animals (Fig. 5A-C). This demonstrated that these cells have the capacity to efficiently migrate from the spleen to the liver (Fig. 5B), with evidence of two nodules present in the bloodstream. Morphologically, the livers contained mainly bile duct-like structures that expressed the cytokeratin markers KRT19 and KRT7, which were easily recognizable after H&E staining (Fig. 5B-C).

**Figure 5:**
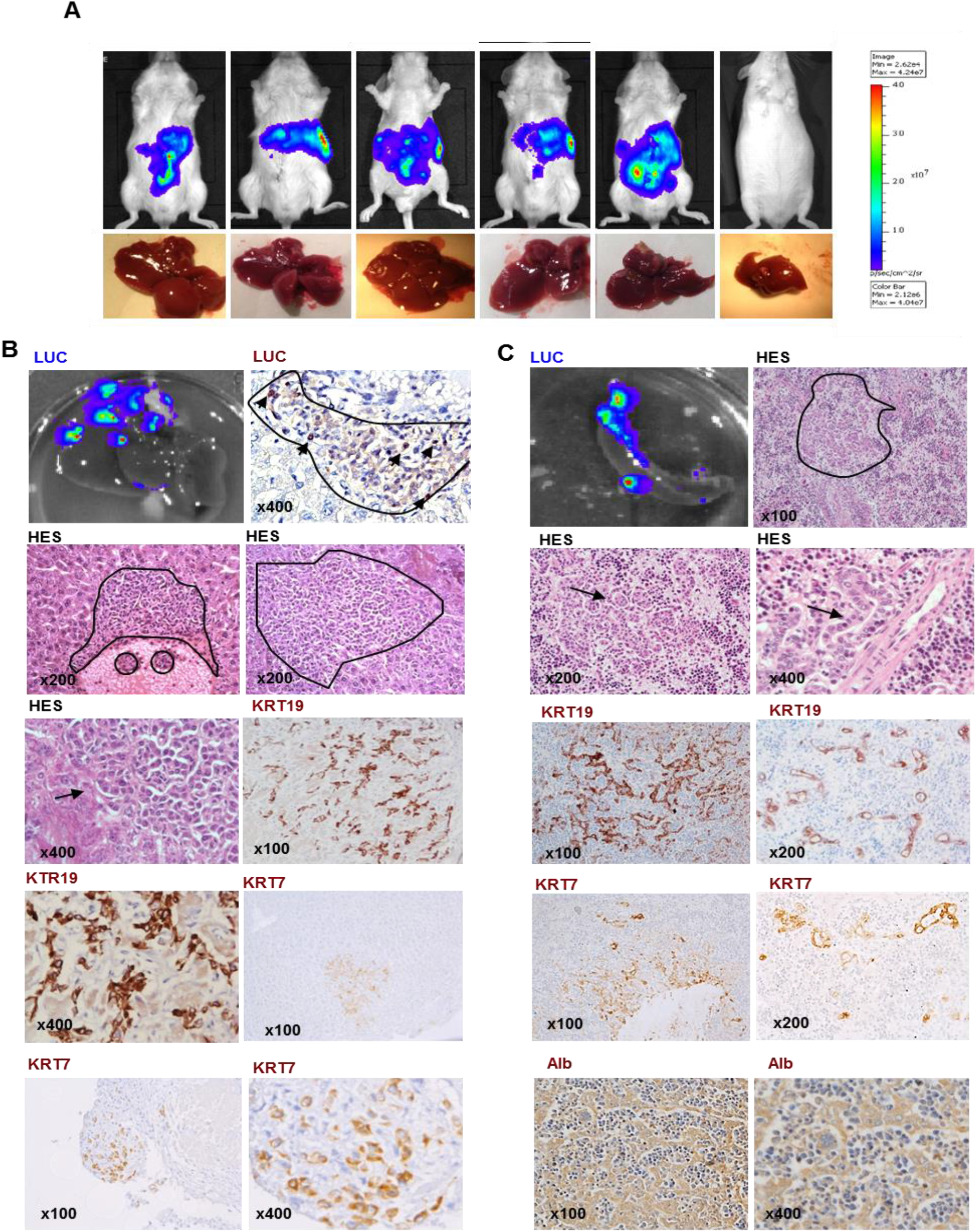
Incorporation of EndoPCs into mouse hepatic parenchyma. **(A)** Bioluminescence imaging for the detection of luciferase gene expression in EndoPCs 7 days after the injection of 3×10^6^ cells in the spleen of five adult C57BL/6 mice that had undergone a partial hepatectomy. PBS was injected into five other mice, which were used as controls. Firefly bioluminescence imaging (Xenogen Ivis 50) was used to detect graft cells. (**B**) Location of luciferase expression in the dissected liver (LUC). Hematoxylin and eosin staining (HES) was used to identify two nodules present in the bloodstream, showing that EndoPCs have the capacity to efficiently migrate from the spleen to the liver. The engrafted tissue shows similarities with liver sinusoid structures after HES and in immunohistochemistry analysis for KRT19 and KRT7. (**C**) Location of luciferase expression in the dissected spleen, showing as many as 8 different sites. HES revealed liver sinusoid structures that resembled biliary ducts, were positive for KRT19 and KRT17, and immunoreactive for Alb.

In addition, EndoPCs were able to survive *in vivo* in the liver microenvironment following orthotropic injection of 3×10^6^ cells of EndoPC-luc in C57BL/6 syngenic adult mice. Luciferase activity increased for as many as 21 days, demonstrating that EndoPCs-luc were able to properly expand in the liver over time (Supporting Fig. 10).

### Differentiation of EndoPCs into hepatocytes and cholangiocytes *in vitro*

We optimized two different differentiation media (DM), which respectively induced the commitment of EndoPCs into either hepatocyte (hDM) or cholangiocyte lineages (cDM) *in vitro*. We first developed a 10-day protocol aimed at directing EndoPCs towards development into hepatocyte-like cells (HLC). These cells expressed high levels of *Alb* mRNA compared to those grown in other culture conditions, and *AFP* transcripts were not detected (Supporting Fig. 11). A significant increase in Alb was observed after differentiation by qRT-PCR (Fig. 6A), ELISA (Fig. 6B), immunofluorescence staining (Fig. 6C), and by flow cytometry analysis, which revealed that 68% of cells were Alb-positive (Fig. 6D).

Differentiated cells possessed typical epithelial morphology and several cells were bi-nucleated (Fig. 6E). Like primary hepatocytes, HLCs accumulated glycogen (Fig. 6E) and showed greater LDL uptake activity than undifferentiated cells did (Fig. 6F). Differentiated hepatocytes also produced lipids, revealed with Oil Red O staining, that were rarer in control conditions (+mLIF) (Fig. 6G). *Ttr, Tat*, cytochrome P450 oxidoreductase (*Por*), and *Cyp3A11* mRNA were also expressed after differentiation (Fig. 6H), and were able to metabolize midazolam to its main metabolite, 1-hydroxymidazolam, by a glucuronidation process. This activity was doubled when a cytochrome P450 inducer—20 μM of pregnenolone-16a-carbonitrile (PCN)—was added (Fig. 6I). Next, we designed a protocol for 3D culture. EndoPCs were seeded directly onto coated Alvetex and differentiated with a two-step protocol that utilized oncostatin M and dexamethasone. After 15 days, 3D-HLCs showed greater Alb expression (Fig. 6J) and LDL uptake activity (Fig. 6K) compared to cells cultivated on 3D scaffolds in culture media without LIF. In addition, the quantity of mRNA transcripts of LIFR dropped down in the HLCs after the hepatogenic commitment of EndoPCs. Finally, after differentiation of the 3D-HLCs, KRT19 was dramatically downregulated, which indicated that the risk of cholangiocytes or biliary duct progenitors growing from these bipotent EndoPCs was low when cultivated on 3D-Alvetex scaffolds (Fig. 6l).

**Figure 6:**
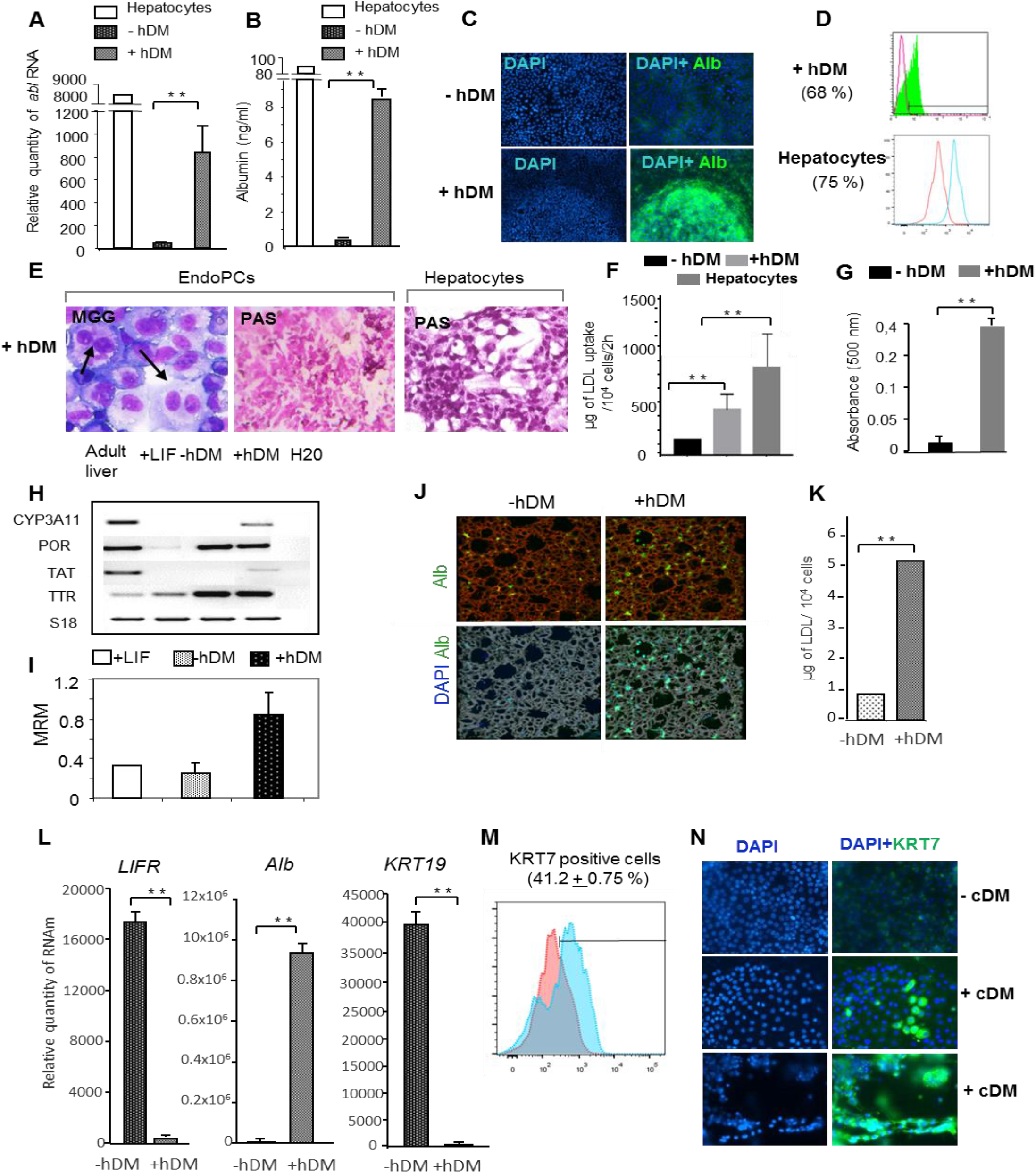
Differentiation of EndoPCs into hepatocyte- and cholangiocyte-like cells. (**A**) Bar graphs showing levels of albumin mRNA as determined by real-time qRT-PCR in EndoPCs before (-hDM) and after differentiation induced by a hepatocyte differentiation medium (+hDM). (**B**) Albumin secretion by murine hepatocytes and by EndoPC-derived hepatocytes after 8 days of culture, as determined by ELISA before (-hDM) and after differentiation (+hDM). (**C**) Immunofluorescence analysis of albumin in EndoPCs before (-hDM) and after differentiation (+hDM). (**D**) Expression of Alb in primary hepatocytes and in EndoPC-derived hepatocytes after differentiation (+hDM), as determined by flow cytometry analysis. (**E**) Morphology of EndoPCs after differentiation (+hDM). Arrow indicates binuclear cells revealed by May-Grünwald-Giemsa (MGG) staining; glycogen was revealed by periodic acid–Schiff (PAS) staining. Primary hepatocytes were used as control. (**F**) Quantification of LDL uptake in murine hepatocytes and in EndoPCs before (-hDM) and after differentiation (+hDM). (**G**) Quantification of lipid content in EndoPCs before (-hDM) and after differentiation (+hDM) using Oil Red O staining. (**H**) Results of RT-PCR used to detect transcripts specific to mature hepatocytes (*Cyp3A11*, *POR*, *TAT*, *TTR*) in EndoPCs cultured with LIF, before (-hDM) and after differentiation (+hDM). (**I**) The pharmacokinetics of midazolam and its main metabolite, 1-hydroxymidazolam, as determined by mass spectrometry of extracts from nondifferentiated EndoPCs (+LIF and without hDM) and differentiated EndoPCs. (**J**) Representative images of EndoPCs that were seeded onto a coated Alvetex 3D culture system, undifferentiated (-hDM) or differentiated (+hDM). Images were taken 15 days after seeding. (**K**) Quantification of LDL uptake of undifferentiated (-hDM) and differentiated (+hDM) EndoPCs after 15 days of culture in an Alvetex 3D culture system. (**L**) Results of RT-PCR used to quantify *LIFR*, *Alb*, and *KRT19* mRNA in Alvetex 3D-derived hepatocytes after 15 days of differentiation, compared to EndoPCs seeded in the Alvetex 3D system but not differentiated (-hDM). (**M**) Expression of KRT7 in EndoPC-derived cholangiocytes as revealed by flow cytometry analysis after 14 days of differentiation, compared to the control (+LIF). (**N**) Immunofluorescence staining for KRT7 in EndoPCs after 14 days of differentiation (+CM) compared to undifferentiated cells (-CM). DNA was stained with DAPI.

To evaluate the quality of hepatocytes derived from EndoPCs, we used Fourier transform infrared (FTIR) spectroscopy, combined with a high brilliance synchrotron radiation source,(*13*) to compare the HLCs to primary adult hepatocytes freshly isolated from the liver of C57BL/6 mice. We compared the spectral signatures using synchrotron infrared micro-spectroscopy and principal component analysis; in this way, we were able to compare the global biochemical composition of the cells, at the level of the individual cell, in terms of their relative content of proteins, polysaccharides, lipids, and nucleic acids. Using this technology, we clearly observed an overlap between the spectra of both types of hepatocytes, which suggested similarity in their metabolisms. Both spectra were distinctly separate from the spectra of non-differentiated EndoPCs, indicating that the biochemical composition of the two groups was quite different (Fig. S12).

We then developed a 14-day cDM protocol to induce the differentiation of EndoPCs into a duct-like phenotype in both 2D and 3sD culture systems, using collagen I as a matrix. The presence of KRT7, which is known to be a specific ductal marker, was assessed using flow cytometry; a mean of 41.2+0.75% of cells were positive for KRT7 compared to the isotype control (Fig. 6M). Histologically differentiated cells displayed a duct-like phenotype: in the 2D culture system, they presented a single-layered non-polarized epithelium, and in the 3D system, bile duct polarized outgrowths, which all expressed KRT7 (Fig. 6N, Fig. S13A). This confirmed the potential of EndoPCs to efficiently differentiate into bile duct-like structures, as occurred spontaneously *in vivo* several weeks after intra-muscular injections (Fig. 4B). We confirmed the duct-like phenotype by performing qRT-PCR, which revealed a dramatic increase in the expression of cholangiocyte markers after differentiation, such as *CFTR*, Gamma-Glutamyltransferase 1 *(GGT1)*, secretin receptor *(SCTR)*, G protein-coupled bile acid receptor 1 *(TGR5), JAG1, NCAM1, SOX9*, and anion exchanger 1 *(AE1)* (Fig. S13B).

## DISCUSSION

LGR5 was recently shown to be a marker of adult stem cells in the murine liver(*4*) that is responsive to both the WNT/β-catenin and LGR5/R-spondin signaling pathways.(*4, 14–17*) To date, this type of cell has been isolated exclusively after inflammation-induced injury *in vivo*(*9, 18–21*) and has been grown *in vitro* only in three-dimensional long-term culture systems that are based on organoids and utilize specific cytokines (*4*).

Our study is the first to demonstrate the feasibility of using terminally differentiated hepatocytes to generate LGR5-positive endodermal progenitor cells (EndoPCs) *in vitro* in 2D long-term culture conditions. Our approach mimicked the epithelial-mesenchymal transition (EMT), and our culturing protocol was unique in that it did not utilize any components from the canonical WNT/β-catenin or LGR5/R-spondin signaling pathways(*4*).

To convert terminally differentiated hepatocytes into hepatocyte-derived progenitors *in vitro*, we used a method that combined transient exposure to the transcriptional factors OCT4, SOX2, KLF4, and c-MYC (OSKM) with STAT3 activators such as IL6 or LIF. These OSKM factors are known to activate an early EMT,(*22*) and STAT3 activators are known to be required for the stimulation of liver regeneration after partial hepatectomy, toxic damage, or inflammation-induced injury(*23*). Our results were consistent with these previous findings, as our EndoPCs expressed markers of mesenchymal stem cells and EndoPC expansion was dependent on LIF or IL6, two closely related cytokines known to bind and activate a common LIF-receptor-gp130 heterodimer, which leads to the phosphorylation of STAT3 by JAK (*24, 25*).

Our protocol was able to efficiently convert terminally differentiated hepatocytes into LGR5^+^ progenitors with stem-like properties, as well as the capacity for self-renewal, clonogenesis, and migration, leading to cells that were resistant to apoptosis. As they were negative for CD117, CD34 (data not shown), and SSEA1, EndoPCs do not represent a novel source of oval cells or iPSCs.

We determined that LGR5^+^ EndoPCs were dependent on three interrelated signaling pathways, gp130/JAK/STAT3, LGR5/R-spondin and WNT/β-catenin which together control their proliferation and capacity for self-renewal. In addition, we found evidence for the existence of cross-talk between the gp130/JAK/STAT3 and WNT/β-catenin signaling pathways and confirmed that IL6 is able to efficiently activate the WNT/β-catenin axis without the use of any other cytokines and/or chemical compounds (*9*).

Importantly, we also found that EndoPCs express the HMG box transcription factor Sox17 and Hepatocyte nuclear factor 4a (HNF4a) in the nucleus. Sox17 is known to be required for definitive endoderm development in mice(*26*), as it interacts with β-catenin, a major component of the WNT signaling pathway, to regulate the transcription of a set of endodermal genes, including Hnf1β, Foxa1 and Foxa2 (*27*). HNF4a is also required for normal liver development, including morphogenesis and differentiation, and controls other transcriptional factors that are critical in liver cell fate as well as genes necessary for hepatocyte function (*28, 29*).

We demonstrated that EndoPCs express both ductal (KRT19) and hepatocyte markers (KRT18), and thus are at least bipotent, with the capacity to commit to either hepatocytes or cholangiocytes *in vitro* and *in vivo*. EndoPCs appeared to retain an epigenetic ‘memory’ of their cell origin, as they readily and efficiently differentiated back into both hepatocytes and cholangiocytes. Our strategy has an advantage over previously published techniques for generating endodermic progenitors, e.g., the work of Zhu and colleagues to directly reprogram fibroblasts into endodermic progenitors. The drawback of their approach is that the differentiation process of the resulting progenitor cells was strongly influenced by the epigenetic memory and the DNA methylation signatures of the fibroblast of origin (*30*). This point was also raised by Shinya Yamanaka, who revealed that reprogrammed iPS cells originating from adult dermal fibroblasts show poor liver differentiation potential (*31*).

In our study, EndoPCs were able to generate macroscopically visible, well-vascularized, and functional liver-like structures after intra-muscular injection, without the use of accessory cells. In this, our results differ from those of Takede (*32*) or Stevens (*33*) and their co-authors, who generated vascularized liver-like buds in mice by mixing endodermal progenitors with human endothelial cells (HUVEC) and mesenchymal stem cells (MCS). In addition, EndoPCs demonstrated the ability to grow and differentiate in a splenic microenvironment, developing into KRT7/19^+^ bile duct-like structures and hepatocytes that secreted albumin. However, even though EndoPCs are bipotent, only a small proportion of undifferentiated cells, and only those with biliary potential, was able to migrate from the spleen to the liver after intra-splenic engraftment, as demonstrated by our finding that only biliary structures were detected in the liver parenchyma. This is consistent with the fact that hepatocytes that have been generated in the spleen can no longer migrate, and instead remain in a relatively quiescent state following terminal differentiation. In agreement with previous studies (*6–8, 34*), our results confirm that hepatocytes are phenotypically plastic, and that, *in vitro*, they are able to change their phenotype through the use of a combination of cytokines to induce an EMT program. By exploiting this technique, it is possible to generate *in vitro* LGR5^+^ liver stem cells with self-renewing capacities. Future research on EndoPCs may be able to develop them into a source for the production of endodermic lineages, therefore opening a vast field of applications in many areas. Furthermore, EndoPCs represent an alternative for the study of physiological liver stem cells and a new source for the large-scale generation of hepatocytes and cholangiocytes for drug screening, toxicity assays, and bioengineered liver approaches.

## MATERIALS AND METHODS

### Establishment of EndoPCs from hepatocytes

Hepatocytes from adult C57BL/6j mice were infected with adenoviruses that encoded OCT4, SOX2, KLF4, and c-MYC. Cells were manually pooled and plated on mitomycin C-arrested mouse embryonic fibroblasts (MEFs) in embryonic stem (ES) cell culture medium that was supplemented with 1000 U/ml of leukemia inhibitory factor (LIF). After 10 to 20 days, colonies were selected and cultured in MEF-free conditions on collagen I-treated plates in ES cell culture medium. After 10 passages the cells were subjected to further analysis. EndoPCs were observed by transmission electron microscopy, as previously described (*35*).

### LGR5 activation and cell culture medium

Cells were cultivated in DMEM/F12 that contained 1% BSA (Sigma) and a cocktail (hereafter referred to as ‘RSPO mix’) of murine cytokines (all from Peprotech): WNT3a (30 ng/ml), RSPO1 (500 ng/ml), EGF (20 ng/ml), HGF (50 ng/ml), and Noggin (50 ng/ml). Organoid-like structures were cultured in the same medium on low attachment plates for 7 days.

### Murine induced pluripotent stem cells

Murine induced pluripotent stem cells (iPSCs) were maintained in culture following the procedure recommended by the manufacturer (iPS02M, ALSTEM/LLC, California).

### EndoPCs FACS analysis and cell immunostaining

Flow cytometry was performed for the detection of SSAEA1, MSC, and oval cell markers using the directly conjugated antibodies anti-SSEA1 (catalogue no. 560142; BD Pharmingen), anti-Sca1 (catalogue no. 553335; BD Pharmingen), anti-CD29 (catalogue no. MCA 2298PET, AbD Serotec), anti-CD51 (catalogue no. 551187; BD Pharmingen), anti-CD166 (catalogue no. 560903; BD Pharmingen), and anti-CD73 (catalogue no. 127205, Biolegend). Standard immunostaining was performed as reported previously.(*36*) Primary antibodies were anti-HNF4α (catalogue no. K9218, Abcam), anti-Foxa2 (catalogue no. AB4125, Millipore), and anti-Sox17 (catalogue no. AF1924, R&D Systems).

### Gene expression profiles of EndoPCs, hepatocytes, and pluripotent stem cells by Plurinet transcriptome analysis

The transcriptomes of EndoPCs were analyzed in triplicate with Affymetrix microarray technology (GeneChip® Mouse Gene 2.0 ST Array). Microarray data were submitted to the public Gene Expression Omnibus (GEO) as GEO dataset GSE51782. The transcriptome dataset from the EndoPCs was compared to transcriptome data in accession GSE33110 (*37*) which includes primary murine hepatocytes, hepatoblasts, embryonic stem cells (D3 and ASE cell lines), hepatoblast-derived iPSCs, and hepatocyte-derived iPSCs (in total 28 samples). The corresponding normalized matrix for each cell type was processed using Illumina Beadchip technology and annotated with the GEO platform GPL6885. Batch cross-normalization was applied to the resulting mixed matrix by decomposition of the variance on experimental batches (*38*). Then, each cross-normalized matrix was dimensionally reduced to the 299 published molecules that make up the Plurinet,(*12*) and an unsupervised principal component analysis was performed on the resulting dataset with the FactoMineR package(*39*).

### Gene expression profiles of EndoPCs, ESCs, and endodermal tissue at stage E8.25 and E11.5

Bioinformatics analyses were performed in the R software environment, version 3.4.3. In order to assess the differentiation stage of EndoPC samples, we downloaded the transcriptome dataset GSE13040 (*40*), which contained a kinetic study of endodermal differentiation, from murine definitive endoderm (E8.25) to specialized endodermal tissues (E11.5). This dataset was processed with Illumina MouseRef-8 v2.0 BeadChip technology, and we annotated the normalized matrix using the GEO platform GPL6885, available on the GEO website. Cross-batch normalization was performed by identifying and removing Fisher variance between types of experiments(*38*). The cross-validated matrix was reduced to the Plurinet gene list,(*12*) and an unsupervised principal component analysis was performed with the R packages FactoMineR(*37*) and factoextra. An expression heatmap was generated with the R bioconductor package made4 (*41*).

### Gene expression profiles of EndoPCs and LGR5^+^ liver stem cells

To identify genes that were differentially expressed between the transcriptomes of sorted LGR5^+^ cells and unsorted hepatocytes, we downloaded the matrix dataset GSE32210 (generated using the Whole Mouse Genome Microarray 4×44K G4122F) from the GEO website. This dataset was first normalized via background subtraction and the Lowess algorithm,(*4*) then annotated with the publicly available transcriptome platform file GPL4134 (https://www.ncbi.nlm.nih.gov/geo/query/acc.cgi?acc=GPL4134). Supervised analysis was performed with the Significance Analysis for Microarray (SAM) algorithm with a false discovery threshold (FDR) of less than 0.05(*42*).

Raw microarray data for EndoPCs were normalized by the RMA method (*43*) and annotated with the custom CDF ENTREZ version 17 (*44*) in the R package bioconductor. Supervised analysis with the Significance Analysis for Microarray (SAM) algorithm was performed to compare EndoPCs and hepatocyte controls. A graphical representation of the gene expression profile and the unsupervised classification was created with the R package gplots using the function heatmap2. Functional enrichment was analyzed with the standalone application Go-Elite version 1.5 using the Wikipathway database (*45*). A functional enrichment score was generated using Cytoscape software, version 3.4.0 (*46*).

### Proliferation assay, cell viability, LDL uptake assay and quantification of CYP3A11 activity

Cells were placed in 24-well tissue culture plates (10^4^ cells per well) and treated with mouse LIF (1000 units/ml, ESG1107, Millipore), 100 ng/ml of murine IL6 (Peprotech, France), or RSPO mix. The gp/JAK/STAT3 pathway was inhibited with 5 μM of pan-JAK inhibitor (CAS 457081-03-7, Millipore). At 24, 48, 72, and 120 hours, Thiazolyl Blue Tetrazolium Bromide (MTT, 5 mg/ml, M2128/Sigma-Aldrich) dye solution was added to each well. A Modulus TM II Microplate reader (Promega) was used to read the absorbance. The viability of the cells was evaluated using 7AAD (BD559925). LDL uptake activity was assessed using the fluorometric LDL uptake assay kit (ab204716). The methodology we used to quantify CYP3A11 activity was derived from that described previously (*47*).

### Western blot

Western blot analysis was performed as previously described(*48*) using anti-STAT3 (9132, Cell Signaling), anti-Phospho-STAT3 (Tyr705) (9131, Cell signaling), anti-LGR5 (AP2745d, Abgent), or anti-β-actin (4970, Cell signaling) as primary antibodies. The blots were incubated for 1 hr with HRP-conjugated secondary antibody (7074, Cell signaling) diluted 1:5000 in 3% BSA/TBS-T. Immunoreactive bands were imaged using an automated platform for the acquisition for chemiluminescence signals (G:BOX Chemi XT4; SYNGENE). For each sample, blotted bands were normalized with respect to β-actin and quantified.

### Intramuscular injection of EndoPCs

All animal studies were approved by the French Ministry of Agriculture. Five million EndoPCs were injected intramuscularly into the flanks of three C57BL/6 mice; several months later, animals were sacrificed and their tissues were prepared for histological analysis using hematoxylin and eosin (H&E) staining. KRT19 and KRT7 were detected by immunohistochemistry staining using anti-KRT19 rabbit antibody (ab52625) and anti-KRT7 rabbit antibody (ab181598). Immuno-reactive albumin was detected using pre-diluted anti-albumin antibody (sc-374670). Glycogen and acid mucosubstances were revealed by staining with periodic acid-Schiff stain (PAS, SD395B) and Alcian blue, respectively.

### Hepatectomy and transplantation in mice

Five C57BL6 mice, 4-8 weeks old (Janvier-Europe, France), were anesthetized by an intraperitoneal injection of ketamine/xylasine. The skin in the upper abdomen and flanks was sterilely prepared and a 2.5-cm median incision was made. A 30% partial hepatectomy of the left and median lobes was performed, and 3×10^6^ EndoPC-luc cells, which expressed luciferase, were injected using a 29G needle into the spleen. In a second experiment, 3×10^6^ EndoPC-luc cells were directly injected into the liver parenchyma of three mice. Bioluminescence imaging was used to monitor graft cells (Xenogen Ivis50).

### Hepatic differentiation of EndoPCs into hepatocytes and cholangiocyte-like cells

To promote the differentiation of EndoPCs into hepatic cells, a two-step protocol was used on collagen-coated plates. EndoPCs were first treated for 5 days with DMEM medium that was supplemented with 15% SVF (ThermoScientific), 1% nonessential amino acids, 1x penicillin-streptomycin, 100 μM β-mercaptoethanol with 1% B27, 25 ng/ml of HGF, and 1 μM of dexamethasone. For the subsequent 5 days, this medium was supplemented with 1% B27, 1 μM of dexamethasone, 20 ng/ml of VEGF, and 20 ng/ml of oncostatin M (OSM). Albumin was quantified in the supernatant with a commercial ELISA kit (Abnova). The methodology used to quantify Cyp3A11 activity was derived from a previously published protocol.(*47*) Quantities of Alb were determined by FACS analysis and by immunocytochemical staining using an anti-albumin antibody (1:100, sc-374670).

EndoPCs were induced to differentiate into cholangiocyte-like cells using a five-step protocol: 1) 3 days of exposure to 25 ng/ml of HGF and 50 ng/ml of EGF; 2) 5 days of exposure to 10 ng/ml of IL6; 3) 2 days of exposure to 10 μM of sodium taurocholate; 4) 2 days of exposure to 1.8 μM sodium butyrate; 5) 2 days of exposure to 20 ng/ml of VEGF. The amount of KRT7 was quantified by FACS analysis using anti-KRT7 murine antibody (1:1500, ab181598).

### Three-dimensional (3D) hepatocyte differentiation system

In order to induce the differentiation of EndoPCs into hepatocyte-like lineages in a 3D-culture system, we used the polystyrene scaffold Alvetex® (200-μm-thick, Reinnervate, Durham, UK) in 6-well format (AVP005-3, 15-mm diameter, Reinnervate). Inserts were coated with MaxGel™ matrix (E0282, Sigma). EndoPCs were plated at a density of 5×10^6^ cells per well in LIF medium for 48 hr. On day 3, cells were subjected to the hepatic differentiation protocol described above. Cells were isolated from the scaffold following the manufacturer’s recommendations and used in an LDL uptake assay and for RNA extraction.

To detect Alb expression and cell distributions in Alvetex, cells were permeabilized in 0.2% TRITON-X (Sigma) for 1 hr at room temperature, then incubated with a mouse polyclonal primary antibody against Alb (1:200; ab106582) overnight at 4°C. Cells were further incubated with an anti-rabbit antibody conjugated to Alexafluor 488 (1:200; Invitrogen; Life Technologies). Images were captured on an inverted spinning disc confocal Leica TCS SP8 microscope with 40× plan objectives. Images were recorded using a color camera and processed using LAS-X software.

### Quantitative real-time polymerase chain reaction (RT-qPCR)

RNA was extracted from the cell pellets using the PureLink RNA Mini Kits (Ambion) according to the manufacturer’s instructions. The mRNA was reverse-transcribed to complementary DNA (cDNA) using the Superscript transcriptase kit (Invitrogen). cDNA was amplified using FastStart SYBR Green Master Mix (Roche). Thermal cycling was performed on an Agilent Mx3000P QPCR System with the following protocol: 1 cycle (50°C/2 min), 1 cycle (95°C/5 min), and 45 cycles (95°C/3 s and 60°C/30s). A melting curve analysis was performed to exclude nonspecific amplification products. Expression values were normalized to those of GAPDH, then relative changes in expression were calculated. Primer sequences are listed in Supplementary Table 4.

### Fourier transform infrared (FTIR) spectroscopy

Cells were cultured directly on low-e slides (Kevley Technologies, OH) for 4 days, then for 12 days in differention medium, then fixed in 10% paraformaldehyde; they were rinsed briefly with distilled water to eliminate salts and dried at room temperature, as described previously (*13*).

### Statistical Analysis

Each experiment was performed at least twice. Statistical significance was evaluated using Student’s t test (unilateral and unpaired).

## Acknowledgments

We would like to express our sincere thanks to Philippe Leclerc, Olivier Feraud, and Dominique Divers for their technical assistance. We would like also to thank Karim Benhioud for the kind gift of PCA350 plasmid, Paule Opolon for anatomopathological analysis, Guillaume Pourcher for *in vivo* experiments, Sylvia Sanquer for metabolic assays and Christophe Sandt for his contribution. Fundings: This work was performed with grants from the ANR “Programme d’Investissements d’Avenir” of the INGESTEM National Infrastructure (ANR-11-INBS-0009-INGESTEM) and INSERM, University Paris Sud.

## Author contributions

D.C: Provision of study material or patients, Data analysis and interpretation, Collection and/or assembly of data, Manuscript writing. C.D: Data analysis and interpretation. N.M.: Provision of study material or patients, Data analysis and interpretation. T.E.: Provision of study material or patients, Data analysis and interpretation. N. O.: Provision of study material or patients, Data analysis and interpretation. J.F.: Conception and design, Final approval of manuscript. A. G.T.: Conception and design, financial support, Administrative support, Final approval of manuscript. A. B-G.:Conception and design, Financial support, Administrative support, Final approval of manuscript. Frank Griscelli: Conception and design, Administrative support, Provision of study material or patients, Collection and/or assembly of data, Data analysis and interpretation, Manuscript writing, Final approval of manuscript.

---

**Supporting Figure 1: Generation of EndoPCs from adult hepatocytes.** (**A**) Timeline of EndoPC generation and diagram of the experimental procedures. Approximately one million hepatocytes, generated from adult mice, were infected with 20 MOI of recombinant adenoviruses that encoded human OCT4, SOX2, cMYC, and KLF4 transgenes. Five days later, cells were manually pooled and plated on mitomycin C-arrested mouse embryonic fibroblasts (MEFs) in embryonic stem (ES) cell culture medium that was changed daily; DMEM medium was supplemented with 1000 U/ml of leukemia inhibitory factor (LIF). As controls, 10^6^ hepatocytes were transduced with an adenovirus that expressed the GFP gene, and 10^6^ non-infected cells were plated on MEFs in the presence of LIF. Murine somatic cells were supplemented with 15% SVF, 1% non-essential amino acids, 1x penicillin-streptomycin, and 100 μM β-mercaptoethanol. After 10 to 20 days, colonies were selected and cultured in MEF-free conditions on collagen I-treated plates in ES cell culture medium. After 10 passages the cells were identified as EndoPCs and further characterized. (**B**) GFP expression, determined by flow cytometry, after transduction of murine hepatocyte cells with 20 MOI of adenovirus that expressed GFP (AdGFP). This dose resulted in the infection of more than 90% of hepatocytes. (**C**) Expression of OCT4, SOX2, cMYC, and KLF4 as determined by Western blot analysis five days after the infection of murine hepatocytes with recombinant adenoviruses. AdGFP-transduced hepatocytes and non-infected hepatocytes were used as controls. (**D**) Morphology of EndoPCs on mitomycin C-arrested MEFs in ES cell culture medium that was supplemented with LIF at passage number 2. (**E**) Morphology of EndoPC clones on a collagen-coated 6-well plate at passage number 5. (**F**) May-Grünwald-Giemsa (MGG) staining of EndoPCs, showing a high nucleus/cytoplasm ratio. (**G**) Transmission electron microscopy (TEM) analysis of EndoPCs, showing cells with a high area of eurochromatin. (**H**) TEM analysis showing abundant lipid droplets (LD) in the cytoplasm.

**Supporting Figure 2:** Karyogram of EndoPCs, showing 40 normal chromosomes. Twenty metaphases were analyzed for each sub-clone and were captured using the Metafer platform (Metasystem version 3.8.6). The karyotype was obtained after G-band staining and the chromosome classification was performed using IKAROS software.

**Supporting Figure 3: Flow cytometry analysis of mesenchymal markers**. Quantification of EndoPCs that were positive for CD73, CD51, CD29, CD166, and Sca1. PE-conjugated anti-mouse antibodies were used to stain 10^5^ EndoPCs, which were then analyzed with a MACS Quant flow cytometer (Miltenyi Biotech, using the MACS Quantify software).

**Supporting Figure 4. Transcriptomic profiles of EndoPCs and different endodermal samples.**

(**A**) Results of a principal component analysis performed on the Plurinet signature of the transcriptome of EndoPC samples, as well as of different endodermal samples from the datasets in GEO accession GSE13040, which included murine embryonic stem cell (D3), definitive endoderm E8.25 (DE), and specialized endodermal samples from liver and pancreas (E11.5). (**B**) Unsupervised clustering of endodermal tissues and pluripotent samples based on the expression patterns of Plurinet genes found in transcriptome analyses. Clustering was based on Euclidean distances.

**Supporting Figure 5**: **EndoPCs express definitive endodermic markers.** (**A**) RT-PCR analysis for the detection of definitive endodermic markers in 15 individual EndoPC clones. (**B**) Immunofluorescence staining showing the expression of Sox17 and Foxa2 in Hepa 1-6 control cells and in EndoPCs. DNA was stained with DAPI.

**Supporting Figure 6: EndoPCs express liver-specific markers.** (**A**) RT-PCR analysis for the detection of markers involved in liver development in eight different EndoPC clones. (**B**) Real-Time Reverse Transcription-PCR for the detection of KRT8/18, KRT7/19, Afp, and Alb in hepatocytes (hep) and EndoPC clones. Target gene expression was normalized relative to that of S18 ribosomal endogenous mRNA. (**C**) Immunofluorescence staining showing the expression of HFN4α-positive cells in Hepa1-6 control cells and in EndoPCs. DNA was stained with DAPI.

**Supporting Figure 7: EndoPCs express pancreatic-specific markers.** (**A**) RT-PCR analysis for the detection of markers involved in endocrine pancreatic development in eight different EndoPC clones. (**B**) Immunofluorescence staining showing the expression of Pdx1 in EndoPCs. DNA was stained with DAPI.

**Supporting Figure 8**: Flow cytometry analysis for the quantification of 7AAD in EndoPCs that expanded up to 168 hours with and without LIF.

**Supporting Figure 9: LGR5/WNT and Jak2/STAT3 pathways regulate EndoPC proliferation.** MTT cell proliferation assay of EndoPCs treated long-term (72 hours of culture) with LIF, IL6, and RSPO mix (1% BSA, 30 ng/ml of mWNT3a, 500 ng/ml of mRspo1, 20 ng/ml of mEGF, 50 ng/ml of mHGF, and 50 ng/ml of mNoggin). The MTT assay revealed a significant increase in proliferation in all treated EndoPCs compared to the non-treated controls. Treating the EndoPCs with 5 μM pan-JAK inhibitor, which inhibited the JAK/STAT3 pathway, resulted in a marked decrease in the proliferation rate among treated EndoPCs (treated with IL6, LIF, and RSPO mix). Tests were performed at least three times and results are presented as the mean ± SD. *P < 0.05, **P < 0.01, ***P < 0.001.

**Supporting Figure 10: LGR5/WNT pathway enhances proliferation potential in EndoPCs treated long-term with RSPO mix**. The MTT proliferation assay revealed a significantly higher proliferation rate in the RSPO-treated EndoPCs compared to their non-treated counterparts after 24, 48, and 72 hrs in culture. Tests were performed at least three times and results are presented as the mean ± SD. *P < 0.05, **P < 0.01, ***P < 0.001.

**Supporting Figure 11:** (**A**) Quantification and (**B**) detection of luciferase activity over a period of 35 days following the injection of 3×10^6^ EndoPCs in the livers of C57BL/6 mice.

**Supporting Figure 12:** Results of RT-PCR used to detect albumin and alpha-fetoprotein (Afp) transcripts in adult hepatocytes, fetal liver cells, and EndoPCs before (+LIF, -hDM) and after differentiation (-LIF, +hDM).

**Supporting Figure 13: Comparison of cell spectra by synchrotron source infrared micro-spectroscopy and principal component (PC) analysis.**

Murine hepatocytes and EndoPCs (before and after differentiation) were compared using infrared microspectroscopy and multivariate statistical analysis. The spectra of single cells were recorded at the SOLEIL synchrotron facility on the SMIS beamline, which exploits the edge and constant field radiations of a bending magnet. Hepatocytes were measured with a dual aperture of 50×50 μm^2^ to match the cell size with the global source. Spectra were recorded with 128-256 scans at 4 cm-1 resolution. Differences between cell lines are put into scale where PC analysis is an unsupervised multivariate statistical analysis that allows the objective comparison of the spectral signatures of over 1500 cells.

**Supporting Figure 14**: Molecular and cellular characterization of EndoPC-derived cholangiocytes.

(**A**) Branching-like structures are shown before (-CM) and after differentiation (+CM) using MGG staining. (**B**) Results of qRT-PCR for the detection of mature cholangiocytes before (-CM) and after differentiation (+CM) of EndoPCs and in Hepa1-6 control cells.

**Supporting Table 1**: Flow cytometry analysis on three different EndoPC clones, showing the percentage of positive cells for each marker analyzed

**Supporting Table 2**: Genes overexpressed in FACS-sorted LGR5^+^ liver cells compared to primary hepatocytes. When the SAM algorithm was used to perform a supervised analysis of the GSE32210 dataset, it identified the overexpression of 337 microarray probes in LGR5^+^ cells (columns describe these genes based on information from different databases and the expression fold-change).

**Supporting Table 3**: LGR5-related genes that were overexpressed in EndoPCs compared to primary hepatocytes. When the SAM algorithm was used to perform a supervised analysis of the EndoPC transcriptome, it identified 88 LGR5-related genes that were overexpressed in EndoPCs compared to primary hepatocytes (columns describe these genes based on information from different databases and the expression fold-change).

**Supporting Table 4:** Primer sequences used for quantitative RT-PCR

