## Supplemental Figures and tables for "Direct reprogramming of adult hepatocytes to generate LGR5+ endodermal progenitor"

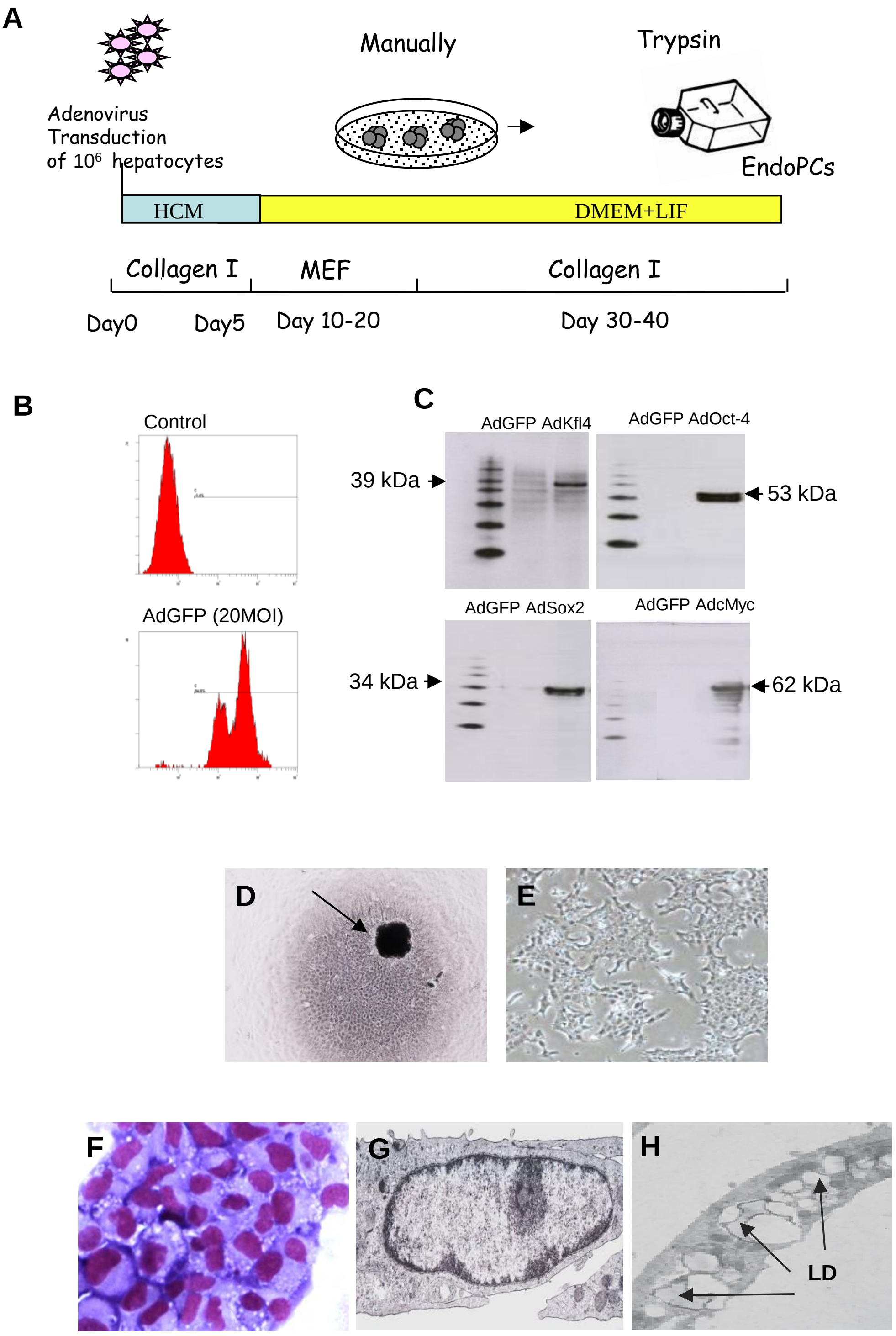


Supporting Figure 1

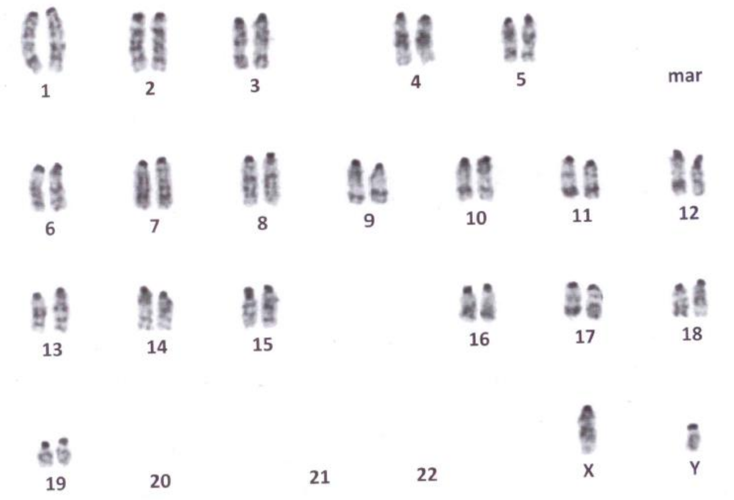


Supporting Figure 2

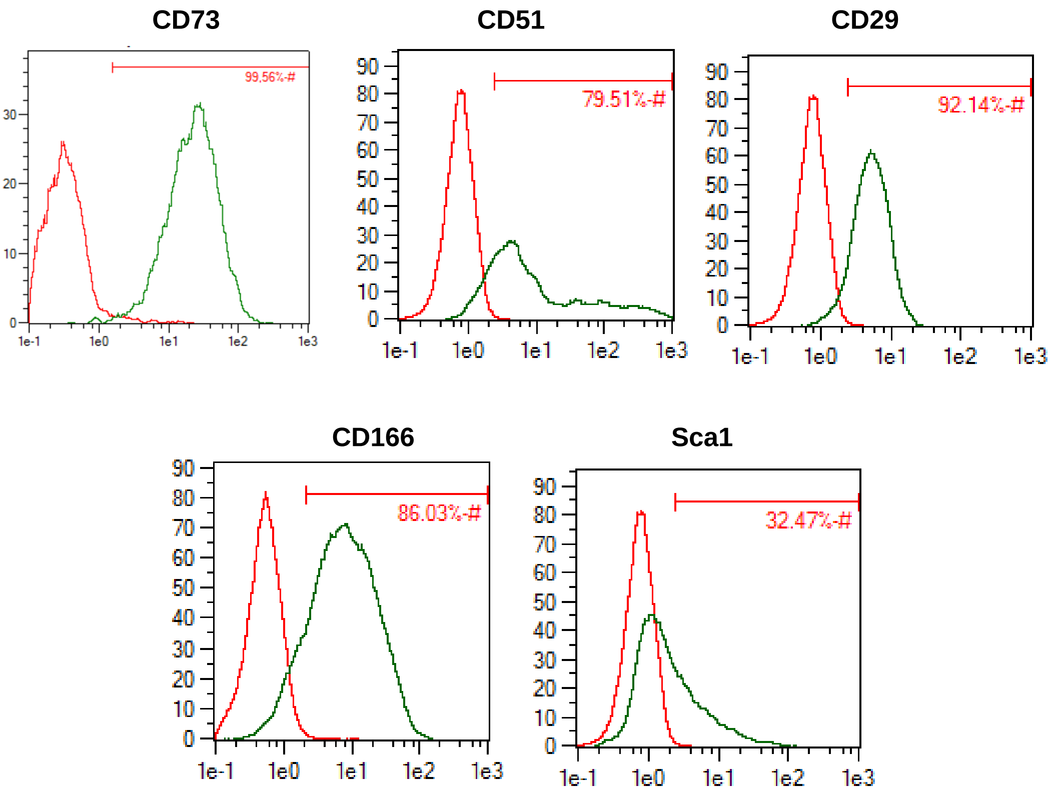


Supporting Figure 3

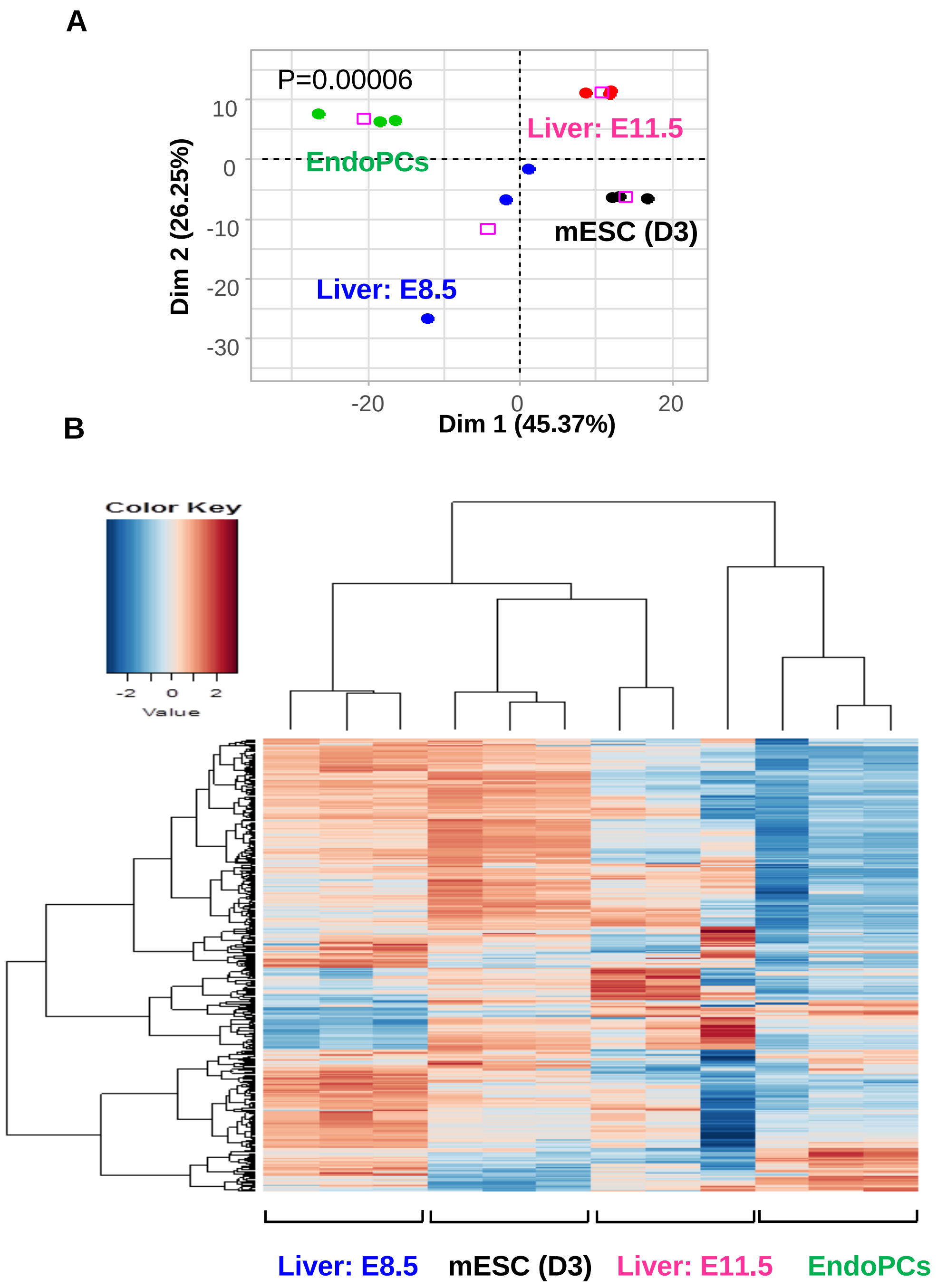


Supporting Figure 4

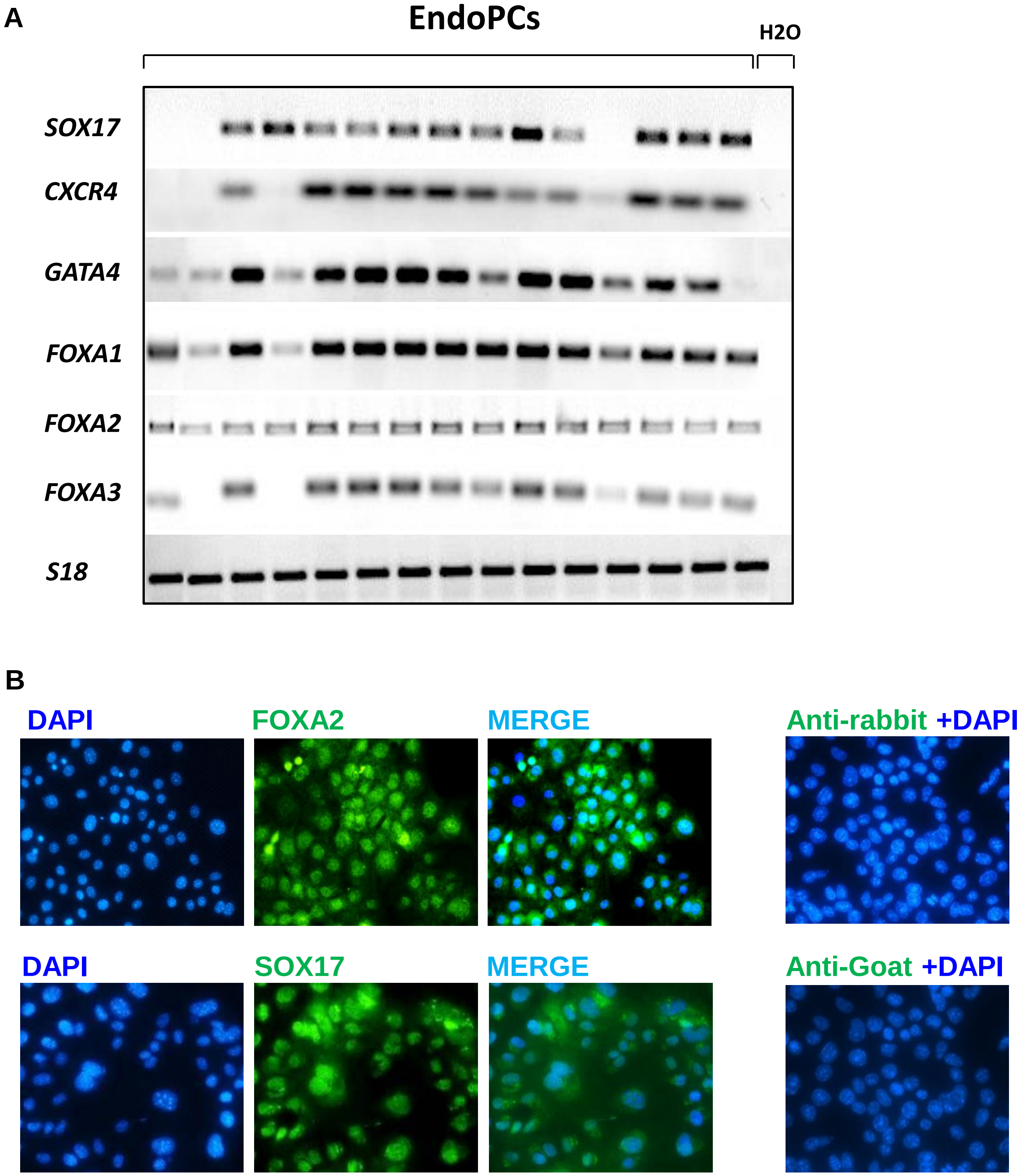


Supporting Figure 5

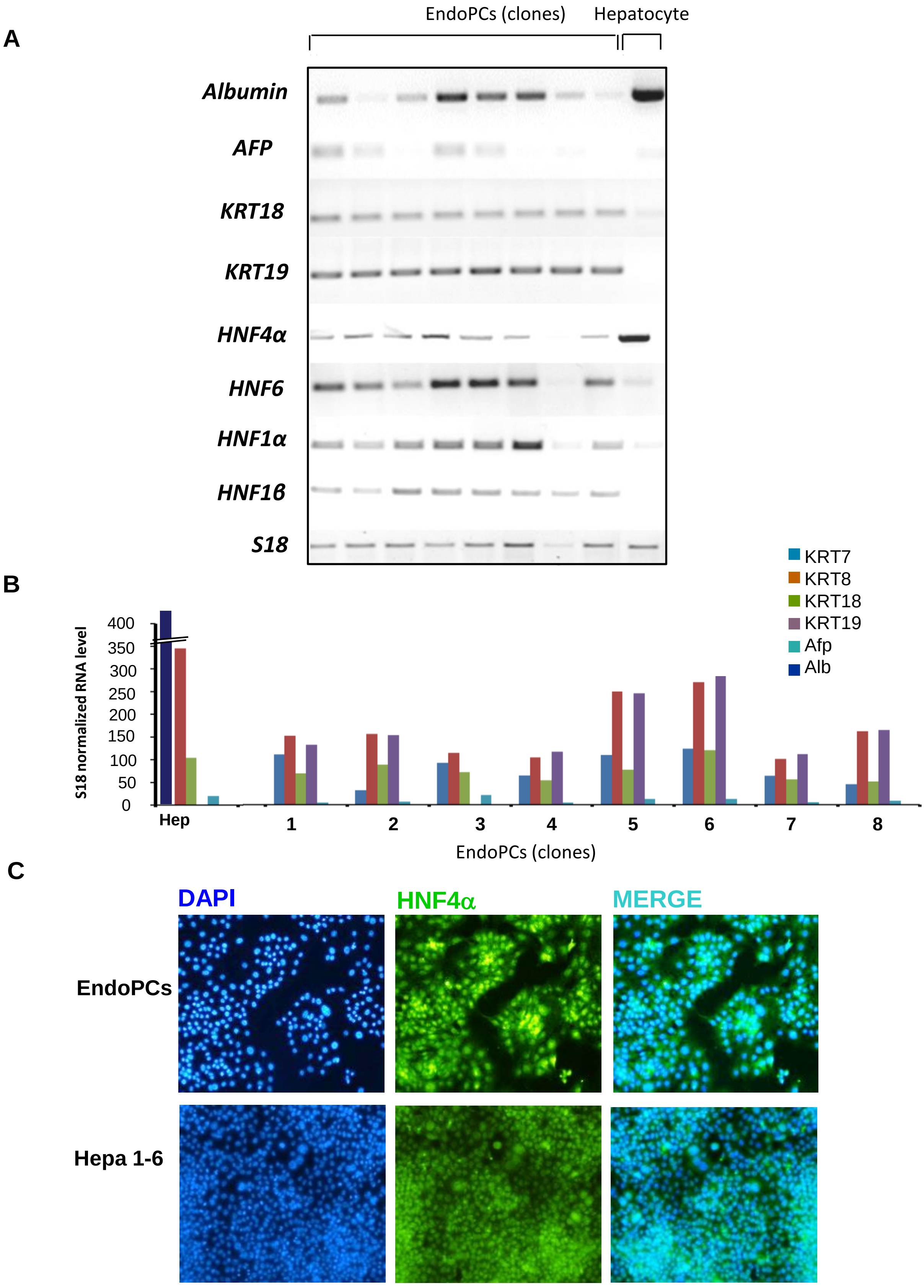


Supporting Figure 6

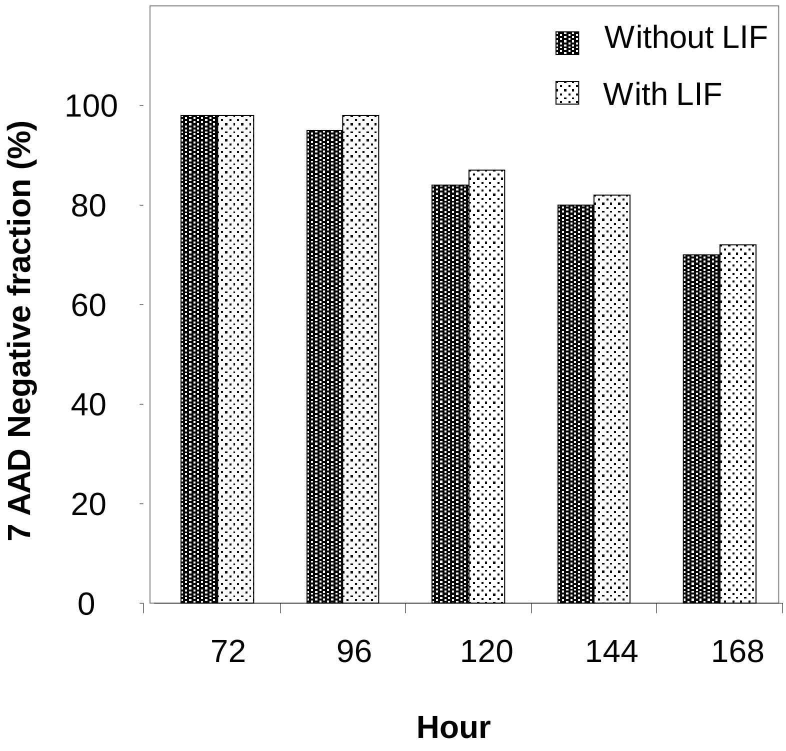


Supporting Figure 7

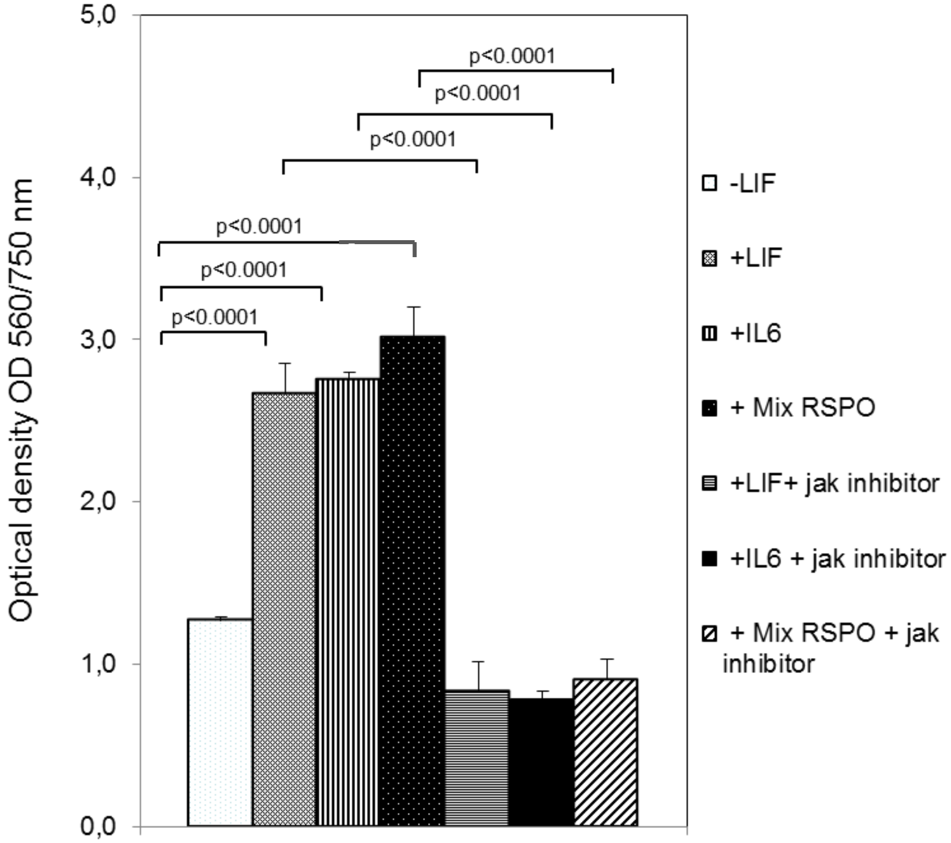


Supporting Figure 8

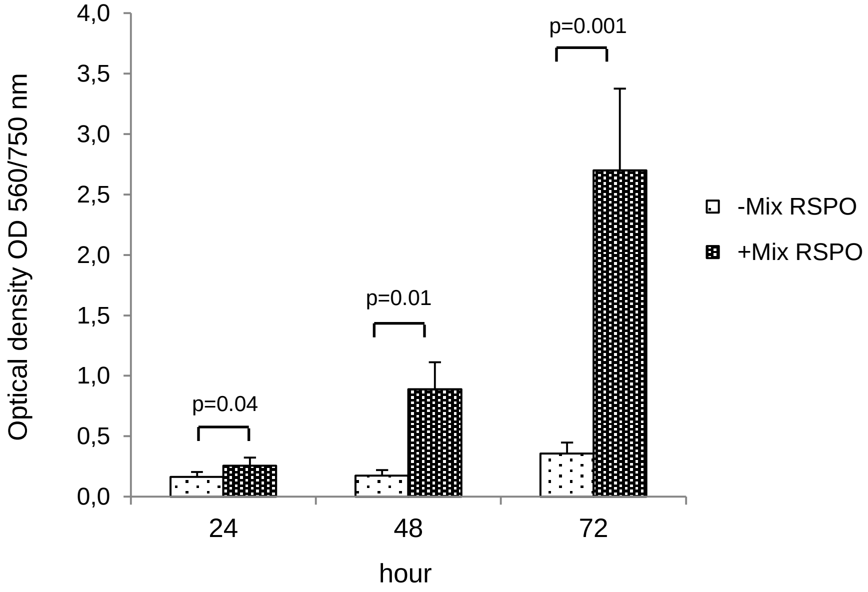


Supporting Figure 9

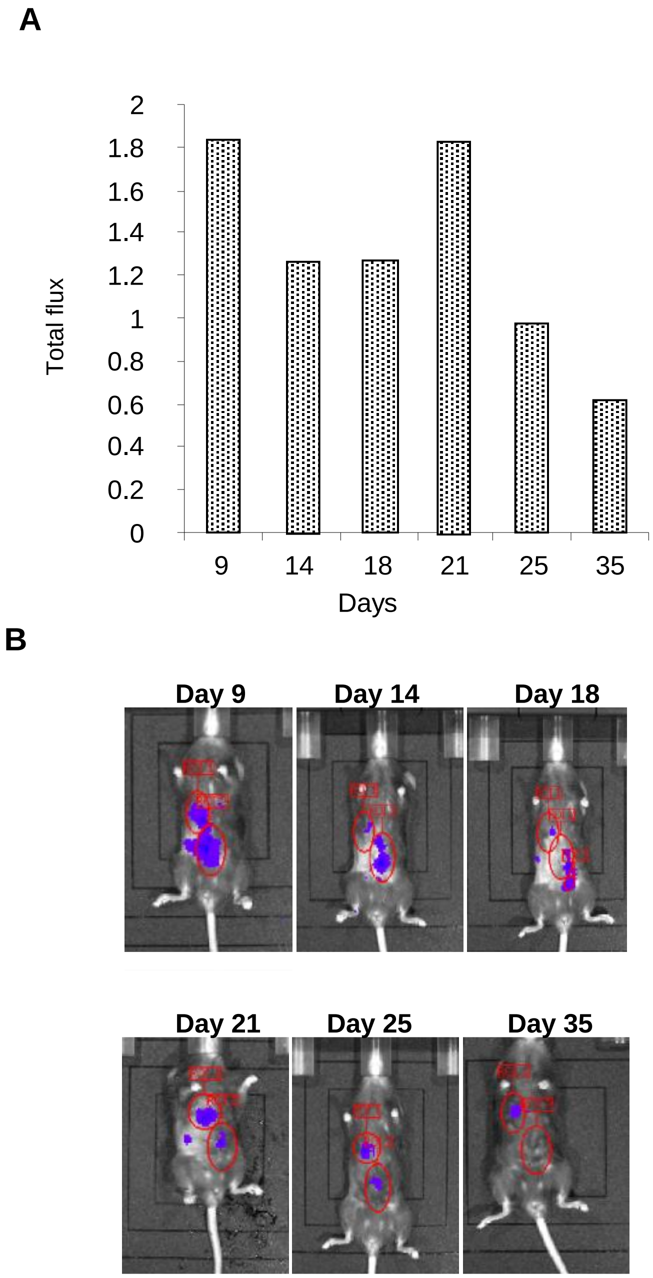


Supporting Figure 10

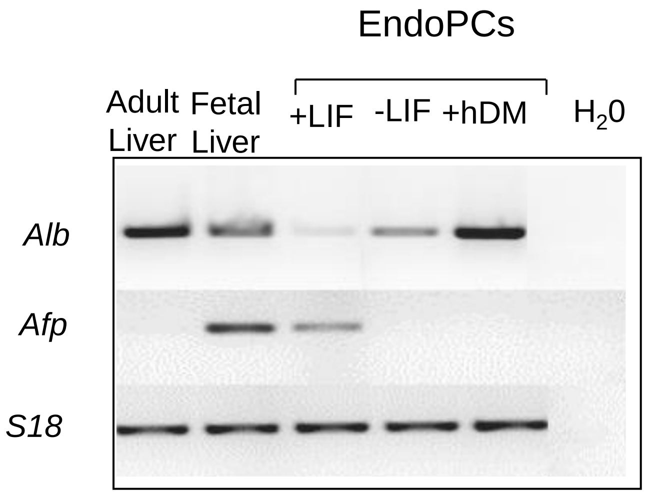


Supporting Figure 11

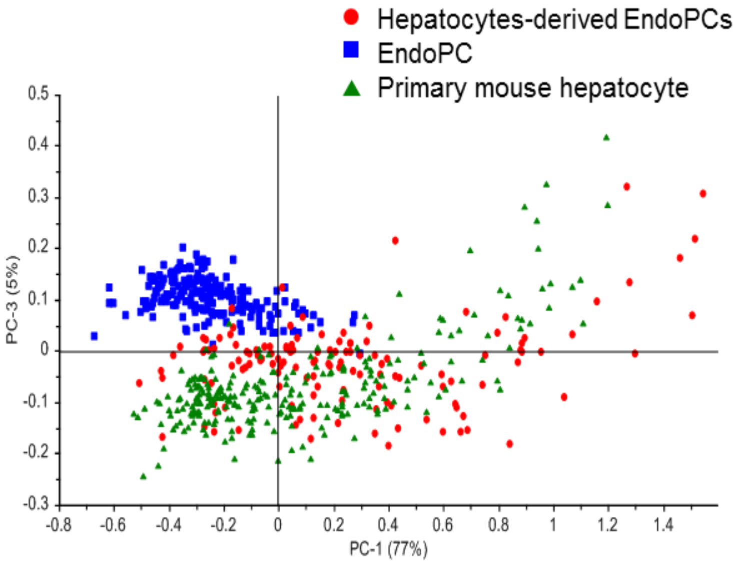


Supporting Figure 12

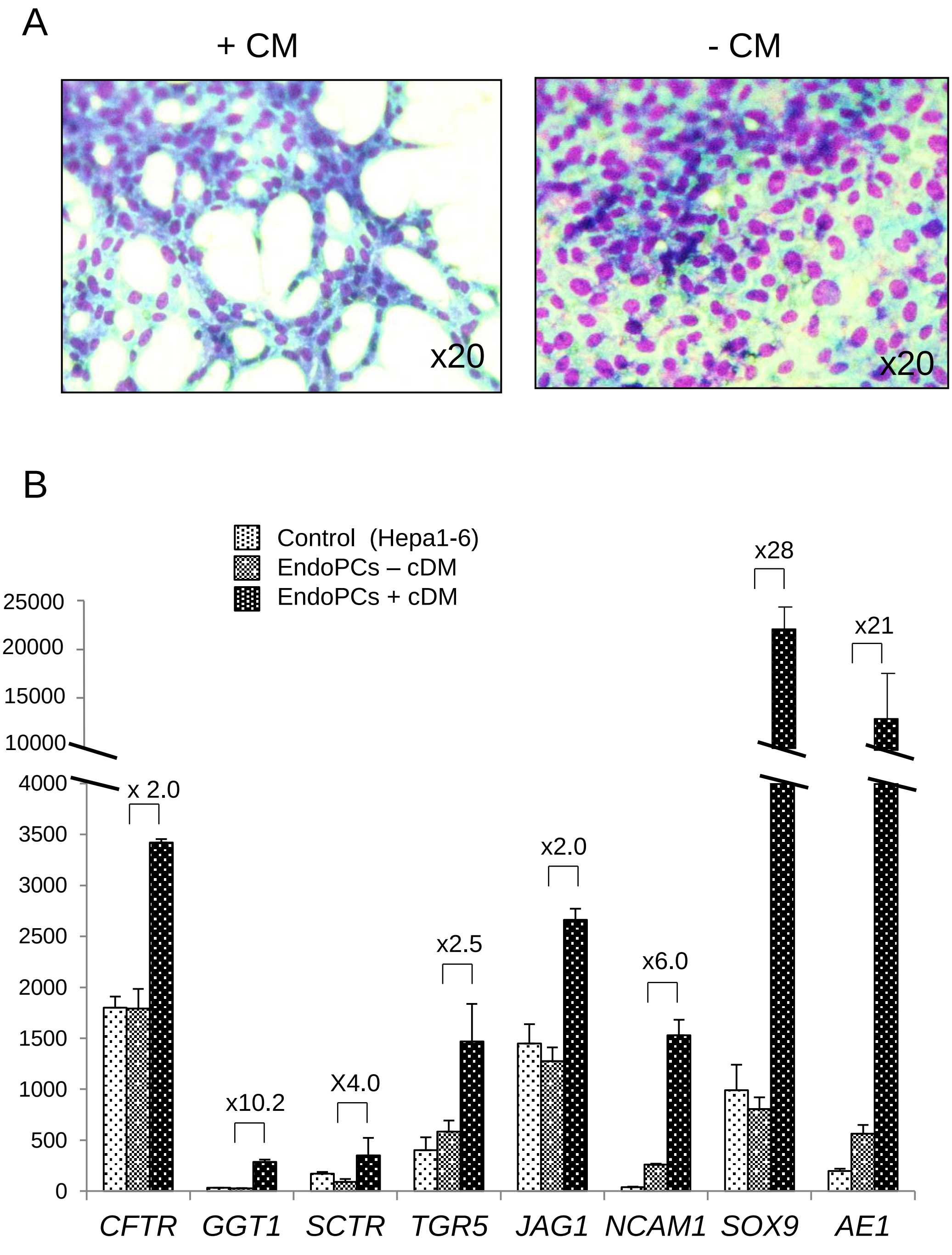


Supporting Figure 13

| | EndoPCs | | |
|---|---|---|---|
| | **#1** | **#2** | **#3** |
| **SSEA1** | 0.22 | 0.20 | 0.18 |
| **CD73** | 98.8 | 96.5 | 88.6 |
| **CD166** | 92.3 | 83.7 | 86.03 |
| **CD51** | 79.5 | 75.5 | 46.23 |
| **CD29** | 92.0 | 85.5 | 70.29 |
| **KRT18** | 97.2 | 97.1 | 86.4 |
| **KRT19** | 96.7 | 98.8 | 90.3 |
| **Sca1** | 32.5 | 30.6 | 25.35 |
| **LGR5** | 83.1 | 99.4 | 94.8 |

Supporting Table1

**Supporting Table 2**

| Gene Symbol | Gene ID | GenBank Accession | Fold change |
|---|---|---|---|
| Mylk | 107589 | AK044527 | 11.12279 |
| 4921530L21Rik | 66732 | NM_025733 | 10.456048 |
| Rfwd2 | 26374 | NM_011931 | 10.39401 |
| Rps27l | 67941 | NM_026467 | 10.143431 |
| Gpt2 | 108682 | NM_173866 | 9.78981 |
| Eef1g | 67160 | AK011951 | 8.826178 |
| Zbtb14 | 22666 | NM_009547 | 8.781705 |
| Slco6d1 | 70866 | AK014872 | 8.75045 |
| Atg4d | 235040 | AK044798 | 8.405915 |
| Zfp36 | 22695 | NM_011756 | 8.039652 |
| Arhgef4 | 226970 | NM_183019 | 8.011314 |
| Hectd3 | 76608 | NM_175244 | 7.7225957 |
| Cdk12 | 69131 | AK137262 | 7.600463 |
| Samt3 | 73495 | NM_028554 | 7.5035214 |
| Gtf2a1 | 83602 | NM_175335 | 7.2962193 |
| Olfr976 | 258364 | NM_146367 | 7.2840195 |
| Vamp1 | 22317 | NM_009496 | 7.219609 |
| St3gal6 | 54613 | NM_018784 | 7.192924 |
| 9430076C15Rik | 320189 | AK035031 | 7.13408 |
| Tmem242 | 70544 | NM_027457 | 7.109999 |
| Sbno2 | 216161 | NM_183426 | 7.1074624 |
| Olfr341 | 258952 | NM_146950 | 7.081101 |
| Clec11a | 20256 | NM_009131 | 7.063682 |
| 4930555G01Rik | 108978 | NM_175393 | 7.000219 |
| Sybu | 319613 | AK051222 | 6.926813 |
| Igf2bp3 | 140488 | NM_023670 | 6.905001 |
| Adarb2 | 94191 | BC059822 | 6.884169 |
| Olfr575 | 259118 | NM_147114 | 6.8309608 |
| Trp53 | 22059 | NM_011640 | 6.796575 |
| Ccdc167 | 68597 | AK003900 | 6.781127 |
| Vmn1r22 | 171196 | NM_134178 | 6.7021904 |
| D3Ertd751e | 73852 | NM_028667 | 6.6864624 |
| Gapdhs | 14447 | NM_008085 | 6.6563315 |
| Mrpl3 | 94062 | NM_053159 | 6.622602 |
| Atp11b | 76295 | NM_029570 | 6.558009 |
| Card11 | 108723 | NM_175362 | 6.40839 |
| Mrps10 | 64657 | NM_183086 | 6.25892 |
| Dfna5 | 54722 | NM_018769 | 6.2239785 |
| Rpl35a | 57808 | AK160963 | 6.209253 |
| Rasa2 | 114713 | NM_053268 | 6.194932 |
| Crb3 | 224912 | NM_177638 | 6.1859603 |
| H2-Oa | 15001 | NM_008206 | 6.1620564 |

| | | | |
|---|---|---|---|
| Zfp37 | 22696 | NM_009554 | 6.1498914 |
| Eif4g1 | 208643 | NM_145941 | 6.1322117 |
| Psrc1 | 56742 | NM_019976 | 6.0071325 |
| Igf1 | 16000 | NM_010512 | 5.9456697 |
| Gm10700 | 1E+08 | AK136892 | 5.927533 |
| Ube3c | 100763 | NM_133907 | 5.904339 |
| Mgat1 | 17308 | NM_010794 | 5.9007883 |
| Pign | 27392 | AK082387 | 5.8987184 |
| Apc | 11789 | AK137301 | 5.8764305 |
| Tmed3 | 66111 | NM_025360 | 5.8564634 |
| Prkg1 | 19091 | AK051624 | 5.826863 |
| Xlr4b | 27083 | NM_021365 | 5.816419 |
| Foxb1 | 64290 | NM_022378 | 5.7879267 |
| Ngfrap1 | 12070 | NM_009750 | 5.7487187 |
| Casp8 | 12370 | NM_009812 | 5.72024 |
| Prl6a1 | 19111 | NM_011166 | 5.668376 |
| Lclat1 | 225010 | NM_001081071 | 5.606109 |
| Camk2d | 108058 | NM_023813 | 5.533771 |
| Taf1a | 21339 | NM_021466 | 5.5145407 |
| Vmn1r9 | 171203 | NM_134185 | 5.494063 |
| Pole2 | 18974 | AK042113 | 5.4620275 |
| Naa20 | 67877 | NM_026425 | 5.454114 |
| Zbtb8a | 73680 | NM_028603 | 5.4539447 |
| Papola | 18789 | NM_011112 | 5.3740783 |
| Fabp4 | 11770 | NM_024406 | 5.354635 |
| LOC102632160 | 1,03E+08 | AK033297 | 5.353326 |
| Fem1b | 14155 | NM_010193 | 5.323753 |
| Tubgcp6 | 328580 | AK052441 | 5.31417 |
| Ankrd27 | 245886 | NM_145633 | 5.31067 |
| Tnni3k | 435766 | AK084817 | 5.29715 |
| Pabpc4 | 230721 | NM_148917 | 5.255092 |
| Slc6a20b | 22599 | NM_011731 | 5.2550316 |
| Tll2 | 24087 | NM_011904 | 5.227288 |
| Dmd | 13405 | NM_007868 | 5.179143 |
| LOC102632051 | 1,03E+08 | AK076935 | 5.167849 |
| Airn | 104103 | AK048015 | 5.1641064 |
| Aldh16a1 | 69748 | NM_145954 | 5.1512604 |
| Shbg | 20415 | NM_011367 | 5.128382 |
| 2810427C15Rik | 69979 | AK013171 | 5.1272664 |
| Mlx | 21428 | NM_011550 | 5.0975246 |
| Pax6 | 18508 | BC036957 | 5.0825667 |
| Nup50 | 18141 | NM_016714 | 5.0281053 |
| Agr3 | 403205 | NM_207531 | 5.0113797 |
| Smg6 | 103677 | NM_001002764 | 4.9909496 |
| Pacrgl | 66768 | NM_025755 | 4.9886513 |
| Ppef2 | 19023 | NM_011148 | 4.98448 |

| | | | |
|---|---|---|---|
| Acn9 | 71238 | AK046436 | 4.961547 |
| Irs1 | 16367 | AY169784 | 4.9577665 |
| Sirt2 | 64383 | NM_022432 | 4.9030337 |
| Clnk | 27278 | NM_013748 | 4.8728204 |
| Foxc2 | 14234 | NM_013519 | 4.870955 |
| Hip1r | 29816 | NM_145070 | 4.8693295 |
| Ccdc115 | 69668 | NM_027159 | 4.859238 |
| Ypel1 | 106369 | NM_023249 | 4.8050175 |
| Mrgprb3 | 404238 | NM_207537 | 4.7801266 |
| Osgin1 | 71839 | NM_027950 | 4.758358 |
| 2610005L07Rik | 381598 | BC086760 | 4.748304 |
| Spty2d1 | 101685 | NM_175318 | 4.7101974 |
| Olfr390 | 258344 | NM_146347 | 4.6966043 |
| 4930519H02Rik | 75058 | AK076692 | 4.6909575 |
| Trak1 | 67095 | BC058971 | 4.682474 |
| Ngrn | 83485 | NM_031375 | 4.679105 |
| Crygd | 12967 | NM_007776 | 4.667646 |
| St8sia6 | 241230 | NM_145838 | 4.6553526 |
| 4930403O15Rik | 73814 | AK015069 | 4.6363525 |
| Gpatch2 | 67769 | NM_026367 | 4.6349173 |
| Bmp2k | 140780 | NM_080708 | 4.6333885 |
| Fbxo28 | 67948 | NM_175127 | 4.625149 |
| Slc5a6 | 330064 | NM_177870 | 4.6222134 |
| Npy2r | 18167 | NM_008731 | 4.6220875 |
| 1700003E16Rik | 71837 | AK005628 | 4.589794 |
| Mtf1 | 17764 | NM_008636 | 4.5759306 |
| Lax1 | 240754 | NM_172842 | 4.572322 |
| A230072C01Rik | 320742 | AK029359 | 4.5579877 |
| Plekha2 | 83436 | NM_031257 | 4.553572 |
| Ccdc176 | 72873 | AK032247 | 4.5284176 |
| Myb | 17863 | NM_010848 | 4.5209937 |
| Vegfa | 22339 | NM_009505 | 4.5132346 |
| Rabggtb | 19352 | BC057661 | 4.5109606 |
| Adhfe1 | 76187 | NM_175236 | 4.493729 |
| Bcl7a | 77045 | NM_029850 | 4.492284 |
| B3gat1 | 76898 | NM_029792 | 4.483408 |
| Bmp2k | 140780 | CO802710 | 4.468369 |
| Abt1 | 30946 | NM_013924 | 4.4661746 |
| Tomm70a | 28185 | NM_138599 | 4.4623437 |
| Mapk3 | 26417 | NM_011952 | 4.4603176 |
| Galnt3 | 14425 | NM_015736 | 4.4125323 |
| Smc2 | 14211 | NM_008017 | 4.410322 |
| Omg | 18377 | NM_019409 | 4.3348017 |
| Tvp23b | 67510 | NM_026210 | 4.328774 |
| Trp53 | 22059 | NM_011640 | 4.3178544 |
| Shisa5 | 66940 | NM_025858 | 4.3045015 |

| | | | |
|---|---|---|---|
| Ubxn7 | 224111 | BC062904 | 4.2939243 |
| 2900006A17Rik | 72913 | AK013487 | 4.2530084 |
| Jsrp1 | 71912 | NM_028001 | 4.2146263 |
| Ranbp3l | 223332 | NM_198024 | 4.209243 |
| Pcdhb17 | 93888 | NM_053142 | 4.2037396 |
| Slc27a4 | 26569 | NM_011989 | 4.18794 |
| Sst | 20604 | NM_009215 | 4.151683 |
| Gpr98 | 110789 | AK081823 | 4.1513643 |
| Fam204a | 76539 | AK044853 | 4.1500626 |
| F630048H11Rik | 1E+08 | AK170335 | 4.1484456 |
| Pacs2 | 217893 | AK122326 | 4.1358986 |
| Wwp2 | 66894 | NM_025830 | 4.1145067 |
| Kntc1 | 208628 | NM_001042421 | 4.1025925 |
| Atp5b | 11947 | NM_016774 | 4.0952024 |
| Wfdc2 | 67701 | NM_026323 | 4.0915227 |
| 4930540M05Rik | 112414 | AK016012 | 4.0831776 |
| Scmh1 | 29871 | AK083889 | 4.080706 |
| Pax2 | 18504 | NM_011037 | 4.0772996 |
| Chsy3 | 78923 | NM_001081328 | 4.052078 |
| Slc25a21 | 217593 | AK044945 | 4.049129 |
| Efna4 | 13639 | NM_007910 | 4.019653 |
| Rftn2 | 74013 | BC038341 | 4.0142965 |
| mars2 | 212679 | NM_175439 | 4.012314 |
| Clasp2 | 76499 | AJ276961 | 4.0069156 |
| Snrnp70 | 20637 | BC049128 | 3.9896255 |
| Vasp | 22323 | NM_009499 | 3.98485 |
| 1110001J03Rik | 66117 | NM_025363 | 3.9818718 |
| Polr3g | 67486 | AK037264 | 3.9771025 |
| Tmem260 | 218989 | NM_172600 | 3.9574082 |
| Klhl2 | 77113 | NM_178633 | 3.9494336 |
| Sys1 | 66460 | NM_025575 | 3.9428585 |
| Phkg1 | 18682 | NM_011079 | 3.9312572 |
| Rell2 | 225392 | NM_153793 | 3.9282918 |
| Gm3877 | 1E+08 | AK080592 | 3.92112 |
| Kcnj6 | 16522 | NM_010606 | 3.92036 |
| Ttc30b | 72421 | AK011097 | 3.9186523 |
| Mars | 216443 | NM_001003913 | 3.9140494 |
| Mib1 | 225164 | NM_144860 | 3.9037557 |
| Cd80 | 12519 | NM_009855 | 3.8976727 |
| Rrad | 56437 | NM_019662 | 3.89504 |
| Plcb2 | 18796 | NM_177568 | 3.8838763 |
| Nipal2 | 223473 | NM_145469 | 3.8806407 |
| Syt13 | 80976 | NM_030725 | 3.8800952 |
| Dhx15 | 13204 | NM_007839 | 3.866127 |
| Ubc | 22190 | NM_019639 | 3.8383827 |
| Fsd1 | 240121 | NM_183178 | 3.8297477 |

| | | | |
|---|---|---|---|
| Nup88 | 19069 | NM_172394 | 3.8290303 |
| Hjurp | 381280 | NM_198652 | 3.8262255 |
| Olfr620 | 258808 | NM_146812 | 3.8073316 |
| Gpr137c | 70713 | NM_027518 | 3.8041728 |
| Slc38a4 | 69354 | NM_027052 | 3.798648 |
| Slc25a42 | 73095 | AK049593 | 3.7941992 |
| Itsn2 | 20403 | NM_011365 | 3.7904477 |
| Pecam1 | 18613 | NM_008816 | 3.7888916 |
| Igf2 | 16002 | NM_010514 | 3.7795153 |
| Il1rap | 16180 | NM_008364 | 3.776532 |
| 1700112J05Rik | 68246 | AK018912 | 3.7653525 |
| Gm12185 | 620913 | BC022776 | 3.7652488 |
| Cic | 71722 | NM_027882 | 3.744131 |
| Reps1 | 19707 | AK042993 | 3.7382588 |
| Mbp | 17196 | NM_010777 | 3.7083793 |
| Chd3 | 216848 | NM_146019 | 3.708205 |
| Ppp1r2-ps7 | 76705 | AK133428 | 3.6903412 |
| Polr3f | 70408 | AK171146 | 3.685033 |
| Gbp7 | 229900 | NM_145545 | 3.682043 |
| Trp53bp1 | 27223 | NM_013735 | 3.6814513 |
| Fabp9 | 21884 | U96149 | 3.669733 |
| Tm9sf1 | 74140 | AK149247 | 3.6439538 |
| Lgalsl | 216551 | NM_173752 | 3.6318376 |
| D030028A08Rik | 319371 | AK050871 | 3.6294997 |
| Porcn | 53627 | NM_023638 | 3.6273923 |
| Zfp664 | 269704 | NM_001081750 | 3.6203725 |
| Ptp4a2 | 19244 | NM_008974 | 3.6202872 |
| Rad51d | 19364 | NM_011235 | 3.6109262 |
| Cyth4 | 72318 | NM_028195 | 3.595358 |
| Uggt2 | 66435 | NM_001081252 | 3.5919564 |
| Vmn1r229 | 171224 | NM_134190 | 3.5714705 |
| Hs3st1 | 15476 | NM_010474 | 3.551712 |
| Asxl3 | 211961 | NM_001167777 | 3.5221162 |
| Sec13 | 110379 | NM_024206 | 3.5181472 |
| Atp8a1 | 11980 | NM_001038999 | 3.5150223 |
| Ska1 | 66468 | NM_025581 | 3.5143807 |
| Usp30 | 100756 | NM_001033202 | 3.4945323 |
| AI413582 | 106672 | NM_001002895 | 3.4828522 |
| Lsr | 54135 | NM_017405 | 3.4733129 |
| Atp13a1 | 170759 | NM_133224 | 3.4660652 |
| Gfap | 14580 | K01347 | 3.464584 |
| Sox7 | 20680 | NM_011446 | 3.4568613 |
| Snapc3 | 77634 | NM_029949 | 3.4538512 |
| Rnf19a | 30945 | NM_013923 | 3.4227085 |
| Wdr90 | 106618 | BC043315 | 3.4164555 |
| Ddx1 | 104721 | NM_134040 | 3.4074156 |

| | | | |
|---|---|---|---|
| Nrarp | 67122 | NM_025980 | 3.399508 |
| BC117090 | 1E+08 | NM_001001332 | 3.3860202 |
| Kcnn1 | 84036 | NM_032397 | 3.3845835 |
| Sptbn1 | 20742 | NM_009260 | 3.3652217 |
| Irak4 | 266632 | NM_029926 | 3.361651 |
| Tmem69 | 230657 | NM_177670 | 3.3511887 |
| Zfp868 | 234362 | AK079745 | 3.3424897 |
| Terf2 | 21750 | NM_009353 | 3.3309846 |
| Men1 | 17283 | NM_008583 | 3.3303554 |
| Tsr1 | 104662 | NM_177325 | 3.3231409 |
| Arhgap24 | 231532 | AK002660 | 3.3132532 |
| Calm2 | 12314 | NM_007589 | 3.2965856 |
| Atad1 | 67979 | NM_026487 | 3.2946699 |
| 0610042G04Rik | 68380 | AK002909 | 3.27517 |
| Amotl1 | 75723 | AK016526 | 3.261146 |
| Atad2 | 70472 | AK037651 | 3.2597084 |
| Agtr1b | 11608 | NM_175086 | 3.2300644 |
| Arx | 11878 | NM_007492 | 3.2285624 |
| Zfp592 | 233410 | NM_178707 | 3.2259815 |
| Blvra | 109778 | NM_026678 | 3.2122946 |
| 4930589P08Rik | 67748 | AK017048 | 3.2069905 |
| Hmga2-ps1 | 15365 | AK033703 | 3.2030365 |
| Unc45b | 217012 | NM_178680 | 3.1903477 |
| Des | 13346 | NM_010043 | 3.178275 |
| Sphk1 | 20698 | NM_025367 | 3.1775534 |
| Tnpo2 | 212999 | NM_145390 | 3.1541429 |
| Spint1 | 20732 | NM_016907 | 3.1481166 |
| 2210016L21Rik | 72357 | NM_028211 | 3.1382368 |
| Ccdc178 | 70950 | NM_027616 | 3.133577 |
| Mapkap1 | 227743 | NM_177345 | 3.1080291 |
| Il12b | 16160 | NM_008352 | 3.1012793 |
| Apitd1 | 69928 | NM_027263 | 3.0936913 |
| Ppp1r9b | 217124 | NM_172261 | 3.0802414 |
| Neb | 17996 | AK086142 | 3.0257704 |
| Il12b | 16160 | NM_008352 | 3.0172136 |
| Rcan1 | 54720 | NM_019466 | 3.0007324 |
| Ubfd1 | 28018 | NM_138589 | 2.9940362 |
| Casd1 | 213819 | BC010201 | 2.9746714 |
| Prss52 | 73382 | NM_028525 | 2.9647543 |
| C130013H08Rik | 1E+08 | AK081392 | 2.9401562 |
| Slc35f5 | 74150 | NM_028787 | 2.9343903 |
| Apobec3 | 80287 | NM_030255 | 2.9285905 |
| Ubc | 22190 | NM_019639 | 2.9202397 |
| Pebp1 | 23980 | NM_018858 | 2.9046772 |
| Scnn1a | 20276 | NM_011324 | 2.8742418 |
| Mx2 | 17858 | NM_013606 | 2.857441 |

Supporting Table 3

| gene ID | mean Hepatocytes | mean EndoPCs | unlog Hepatocytes | unlog EndoPcs | Fold change EndoPCs/hepatocytes |
|---|---|---|---|---|---|
| Ngfrap1 | 6.40 | 11.21 | 84.54 | 2378.33 | 28.13 |
| Galnt3 | 4.97 | 9.27 | 31.39 | 620.04 | 19.75 |
| Pcdhb17 | 3.46 | 6.58 | 11.03 | 96.15 | 8.72 |
| Pax2 | 4.27 | 7.37 | 19.29 | 166.21 | 8.61 |
| Atad2 | 5.39 | 8.44 | 42.02 | 347.53 | 8.27 |
| Smc2 | 5.54 | 8.57 | 46.45 | 380.34 | 8.19 |
| Kntc1 | 4.46 | 7.45 | 21.97 | 175.25 | 7.98 |
| Pole2 | 3.63 | 6.62 | 12.41 | 98.56 | 7.94 |
| Plekha2 | 4.33 | 6.95 | 20.15 | 123.54 | 6.13 |
| Fem1b | 6.31 | 8.73 | 79.46 | 423.18 | 5.33 |
| Trp53 | 6.57 | 8.87 | 95.10 | 466.67 | 4.91 |
| Ska1 | 4.82 | 7.02 | 28.26 | 129.38 | 4.58 |
| Psrc1 | 5.20 | 7.30 | 36.84 | 157.37 | 4.27 |
| Eef1g | 8.40 | 10.29 | 338.90 | 1249.27 | 3.69 |
| Rasa2 | 5.42 | 7.26 | 42.90 | 153.51 | 3.58 |
| Igf2 | 5.71 | 7.44 | 52.42 | 173.64 | 3.31 |
| Zfp37 | 4.00 | 5.73 | 16.04 | 53.05 | 3.31 |
| Ppp1r9b | 4.86 | 6.55 | 29.06 | 93.86 | 3.23 |
| Hs3st1 | 4.20 | 5.88 | 18.39 | 58.76 | 3.20 |
| Uggt2 | 4.32 | 5.95 | 19.92 | 61.91 | 3.11 |
| Camk2d | 5.43 | 7.02 | 43.22 | 129.35 | 2.99 |
| Mapk3 | 8.07 | 9.65 | 269.12 | 803.35 | 2.99 |
| Pign | 6.20 | 7.77 | 73.44 | 217.79 | 2.97 |
| Efna4 | 4.62 | 6.08 | 24.61 | 67.46 | 2.74 |
| Rabggtb | 5.25 | 6.68 | 37.96 | 102.88 | 2.71 |
| Hmga2-ps1 | 4.54 | 5.94 | 23.29 | 61.37 | 2.64 |
| Fbxo28 | 6.28 | 7.68 | 77.83 | 205.05 | 2.63 |
| Zfp664 | 5.63 | 7.00 | 49.56 | 127.94 | 2.58 |
| Pacs2 | 6.53 | 7.84 | 92.53 | 229.03 | 2.48 |
| Snrnp70 | 8.70 | 9.94 | 415.21 | 979.63 | 2.36 |
| Naa20 | 6.83 | 8.07 | 113.66 | 267.98 | 2.36 |
| Zfp868 | 4.86 | 6.04 | 29.10 | 65.99 | 2.27 |
| Amotl1 | 5.51 | 6.68 | 45.50 | 102.71 | 2.26 |
| Cic | 6.66 | 7.81 | 101.26 | 225.17 | 2.22 |
| Tubgcp6 | 5.41 | 6.55 | 42.55 | 93.42 | 2.20 |
| Clasp2 | 6.03 | 7.14 | 65.32 | 140.70 | 2.15 |
| Snapc3 | 6.27 | 7.37 | 77.04 | 165.51 | 2.15 |
| Apc | 6.42 | 7.52 | 85.81 | 183.10 | 2.13 |
| Men1 | 7.07 | 8.16 | 134.48 | 285.48 | 2.12 |
| Cdk12 | 6.97 | 8.04 | 125.51 | 264.01 | 2.10 |
| Dhx15 | 8.99 | 10.05 | 508.31 | 1063.00 | 2.09 |
| Pebp1 | 5.86 | 6.89 | 57.93 | 119.00 | 2.05 |

| | | | | | |
|---|---|---|---|---|---|
| Chd3 | 10.70 | 11.73 | 1657.89 | 3385.38 | 2.04 |
| Terf2 | 5.92 | 6.95 | 60.72 | 123.32 | 2.03 |
| Calm2 | 11.03 | 12.04 | 2095.87 | 4215.92 | 2.01 |
| Trp53bp1 | 5.85 | 6.84 | 57.85 | 114.84 | 1.99 |
| Mbp | 4.36 | 5.35 | 20.54 | 40.66 | 1.98 |
| Tnpo2 | 7.48 | 8.44 | 178.01 | 348.35 | 1.96 |
| Sbno2 | 6.12 | 7.07 | 69.47 | 134.07 | 1.93 |
| Porcn | 6.09 | 7.01 | 68.09 | 128.70 | 1.89 |
| Bmp2k | 6.45 | 7.36 | 87.15 | 163.96 | 1.88 |
| Wdr90 | 4.92 | 5.80 | 30.23 | 55.84 | 1.85 |
| Atp13a1 | 6.61 | 7.49 | 97.63 | 179.23 | 1.84 |
| Hjurp | 8.11 | 8.97 | 276.76 | 500.16 | 1.81 |
| Omg | 4.45 | 5.29 | 21.79 | 39.01 | 1.79 |
| Nup88 | 7.40 | 8.23 | 168.47 | 299.32 | 1.78 |
| Atad1 | 9.06 | 9.88 | 533.26 | 942.08 | 1.77 |
| Rfwd2 | 8.90 | 9.71 | 476.79 | 835.21 | 1.75 |
| Tsr1 | 7.39 | 8.20 | 168.13 | 293.52 | 1.75 |
| Airn | 4.38 | 5.17 | 20.84 | 36.12 | 1.73 |
| Casp8 | 7.51 | 8.29 | 181.77 | 313.60 | 1.73 |
| Bcl7a | 6.24 | 7.02 | 75.69 | 130.11 | 1.72 |
| AI413582 | 5.95 | 6.73 | 61.85 | 106.07 | 1.71 |
| Ddx1 | 9.29 | 10.07 | 627.62 | 1072.16 | 1.71 |
| Polr3f | 5.54 | 6.31 | 46.57 | 79.20 | 1.70 |
| Nup50 | 7.01 | 7.77 | 128.51 | 218.51 | 1.70 |
| St3gal6 | 4.52 | 5.28 | 22.97 | 38.76 | 1.69 |
| Shisa5 | 5.33 | 6.08 | 40.35 | 67.49 | 1.67 |
| Ypel1 | 5.31 | 6.03 | 39.76 | 65.50 | 1.65 |
| Arhgap24 | 5.22 | 5.93 | 37.23 | 61.15 | 1.64 |
| Vasp | 5.76 | 6.46 | 54.29 | 87.99 | 1.62 |
| Rrad | 4.98 | 5.68 | 31.61 | 51.15 | 1.62 |
| Irs1 | 6.38 | 7.04 | 83.07 | 131.61 | 1.58 |
| Wwp2 | 6.56 | 7.21 | 94.28 | 148.32 | 1.57 |
| Apitd1 | 5.06 | 5.67 | 33.28 | 51.05 | 1.53 |
| Ubxn7 | 7.35 | 7.97 | 163.54 | 250.43 | 1.53 |
| Pacrgl | 6.75 | 7.34 | 107.34 | 162.30 | 1.51 |
| Mib1 | 8.10 | 8.68 | 274.07 | 410.92 | 1.50 |
| Sphk1 | 4.76 | 5.34 | 27.18 | 40.45 | 1.49 |
| Papola | 8.24 | 8.76 | 301.94 | 432.31 | 1.43 |
| Usp30 | 6.77 | 7.29 | 109.24 | 156.25 | 1.43 |
| Gtf2a1 | 8.31 | 8.80 | 316.48 | 446.44 | 1.41 |
| Taf1a | 5.67 | 6.16 | 50.99 | 71.27 | 1.40 |
| Eif4g1 | 9.46 | 9.94 | 705.20 | 980.84 | 1.39 |
| Itsn2 | 7.25 | 7.72 | 152.28 | 211.09 | 1.39 |
| Smg6 | 6.26 | 6.73 | 76.87 | 106.06 | 1.38 |
| Des | 4.34 | 4.77 | 20.21 | 27.35 | 1.35 |
| Ube3c | 7.66 | 8.08 | 202.88 | 270.39 | 1.33 |

### Supporting Table 4

| Gene | Forward Primer 5'- 3' | Reverse Primer 5'- 3' |
|---|---|---|
| AE1 | CTGGAGGTGGATAGAGAGCG | CTCCGCATTCTCAGGGATCT |
| AFP | GACTGCTCGAAACATCCCAC | TTGGACGCAGCGAAATGTAG |
| ALB | AACCTAGGAAGAGTGGGCAC | AGACACACACGGTTCAGGAT |
| ALP | GGACAGGACACACACACACA | CAAACAGGAGAGCCACTTCA |
| AQP1 | GACCCTTCCCCTTGAACTCA | CCCTTCTAGAACCTGTGGCA |
| AXIN2 | AAGAAGGAGACCGGTCACAG | GGTCCTGGGTAAATGGGTGA |
| CFTR | GTCGTCTCGGCATTACAACC | CCAGTTGTTTGAGCTGCTGT |
| C-MYC | CTGTCCATTCAAGCAGACGA | TCCAGCTCCTCCTCGAGTTA |
| CXCR4 | GCTGGCTGAAAAGGCAGTCTAT | TGACGTCGGCAAAGATGAAGT |
| CYP2C55 | TCAAAAGTCTATGGCCCCGT | TCCCCATCCCAAAACTCCTC |
| CYP2D22 | CCTCATCACCAACCTGTCCT | CAGGAGGCAGGTGAAGAAGA |
| CYP2E1 | TTGAAGCCTCTCGTTGACCC | CGTGGTGGGATACAGCAA |
| CYP3A11 | GACAAACAAGCAGGATGG | AATGTGGGGGACAGCAAAG |
| FOXA1 | ACCCCACGAATCTCAGCTGCA | TCTCCTTATGGCGCTACCTTG |
| FOXA2 | CCATCCAGCAGAGCCCCAACA | TTCTGGACCCTGCACCCAGAC |
| FOXA3 | TCGTCCACACCTTATTTCAGCG | GCTCTCTGCTTAATGCATCCT |
| G6P | CAGGACTGGTTCATCCTT | ACCGACTACTACAGCAAC |
| GAPDH | GAGAGGCCCTATCCCAACTC | TCAAGAGAGTAGGGAGGGCT |
| GATA4 | GAGCCTGCCAAGCCAAGC | GGACAAAGGTGATAGACGGGAG |
| GGT1 | TAGCGACCCCTGAACAGAAG | CATGTTGCGGATCACCTGAG |
| GLUT2 | CTCTGAAGACGCCAGGAATTCCAT | CGGTGGGACTTGTGCTGCTGG |
| IAPP | CTGTGGCACTGAACCACTTG | GGTTCGTTCCAGCAACAACC |
| INS1 | CCTGTTGGTGCACTTCCTAC | TGCAGTAGTTCTCCAGCTGG |
| INS2 | GCAGCACCTTTGTGGTTCCC | TGCAGTAGTTCTCCAGCTGG |
| JAG1 | GTGGACGGAGACAACTGGTA | CAAAGGCACAAGGGGAAGAC |
| KLF4 | AGGGTCTGCTACTGAGATGCTCTG | TTAGGCTGTCGGGGCCACGA |
| KRT18 | CGATACAAGGCACAGATGGA | CTTCTCCATCCTCCAGCAAG |
| KRT19 | ATTGAGGAGCTGAACACCCA | CTCAATCTCAAGGCCCTGGA |
| KRT7 | TTCCCCGAATCTTTGAGGCT | TCTTCCACCACATCCTGCAT |
| KRT8 | ATCGAGATCACCACCTACCG | TGAAGCCAGGGCTAGTGAGT |
| LGR5 | AACGGTCCTGTGAGTCAACC | AGTCATGGGGTAAGCTGGTG |
| LIFR | AACAGCAAGGACAATGGCAG | CCCCTCTCTTCAGACCGTTT |
| LIN28 | TCCTGCACTGTGTTCTCAGG | GCACTCCAAATGGTTGGTCT |
| NANOG | CAACCACTGGTTTTTCTGCCACCG | AGGGTCTGCTACTGAGATGCTCTG |
| NCAM1 | TGTGTCAAGTGGCAGGAGAT | TGAGGGTAGAGGAGTCGTCA |
| NCAM2 | GGTGTCCCCTCAAGAGTTCA | GGATGGTGGTGACTTCCTCA |
| NEUROD1 | GATGCGAATGGCTATCGAAAG | CTGATCTGGTCTCCTTCGTACAG |
| NGN3 | TGGCACTCAGCAAACAGCGA | AGATGCTTGAGAGCCTCCAC |
| NKX2.2 | AACAACCGTGGTAAGGATCG | CTCTTCTCCAAAGCGCAGAC |
| NKX6.1 | TTCTCTGGACAGCAAATCTTCG | CTGAGTGATTTTCTCGTCGTCA |
| Oct3-4 | CCTTGCAGCTCAGCCTTAAG | GCGATGTGAGTGATCTGCTG |
| PDX1 | GGTGCTTACACAGCGGAACC | CCTACTGCCTTCGGGCCTTA |
| POR | TAGTCTTCTGCATGGCCACA | TGGTCCACATACTTGCCCAT |
| S18 | AGGAATTGACGGAAGGGCAC | GTGCAGCCCCGGACATCTAAG |
| SCTR | TTTGTTGCTTTGTGGGCTGT | AATCACAGGCCCTCGAATGA |
| SOX17 | GCCAAAGACGAACGCAAGCGGT | TCATGCGCTTCACCTGCTTG |
| SOX2 | GGAGTGGAAACTTTTGTCCGA | TTCATGTTAGGTCTGCGAGCTG |
| SOX9 | AGATAAGTTCCCCGTGTGCA | TGACGTGTGGCTTGTTCTTG |
| STAT3 | CGAGAGCAGCAAAGAAGGAG | TGGTCGCATCCATGATCTTA |
| TAT | TGGCGGCCGCAACCCTTCAGTGGGT | TGGACTAGTGCTTGGGCTTGAGATTAC |
| TCF4 | CTCACGCCTCTCATCACGTA | TCCTGTCGTGATTGGGTACA |
| TGR5 | CTCATCTCATTGGGCAGCAC | GGATTGTCCCTCTTGGCTCT |
| TTR | TGCTGTAGACGTGGCTGTAA | ACCACATCCGCGAATTCATG |
| BCATENIN | AGGGTGCTATTCCACGACTA | CACCCTTCTACTATCTCCTCCAT |
